# Oncogenic HPV types identified in Paleolithic and Chalcolithic human genome sequencing data from Ust’-Ishim and Ötzi

**DOI:** 10.64898/2025.12.14.694221

**Authors:** Juliana B. Yazigi, Caio O. Cyrino, Cristina M. Peter, Renata C. Ferreira, Juliana T. Maricato, Luiz M. Janini, Marcelo R.S. Briones

**Affiliations:** Center for Medical Bioinformatics, Escola Paulista de Medicina, Federal University of São Paulo (UNIFESP), São Paulo 04039-032, SP, Brazil; Departamento de Microbiologia, Imunologia e Parasitologia, Escola Paulista de Medicina, Federal University of São Paulo (UNIFESP), São Paulo 04023-062, SP, Brazil; Graduate Program in Bioinformatics, University of São Paulo, São Paulo 05508-000, SP, Brazil; Epigene LLC, São Paulo 01449-070, SP, Brazil

**Keywords:** Paleovirology, Ancient DNA, HPV, Oncogenesis, Human evolution

## Abstract

Human papillomaviruses (HPVs) are ancient viruses with diverse lineages infecting epithelial tissues in primates and humans. Although contemporary distribution and clinical importance are well understood, there is limited knowledge about their occurrence among prehistoric human populations. We investigated the presence of HPV in ancient anatomically modern humans (AMHs) by analyzing genome sequencing data from two exceptionally preserved individuals: Ust’-Ishim (∼45,000 years BP) and Ötzi the Iceman (∼5,300 years BP). Using a combination of reference-guided mapping and ancient DNA authentication criteria, we searched for HPV sequences in these ancient genomes. We detected high-confidence papillomavirus fragments in both individuals. Further phylogenetic and comparative analyses revealed that the reconstructed sequences belong to HPV16, the most oncogenic HPV lineage. Our study presents the earliest molecular evidence of HPV16 in anatomically modern humans (AMHs), pushing back its evolutionary history and challenging the idea that HPV16A entered *Homo sapiens* through Neanderthal interbreeding. Our results suggest that HPV16 was already present in modern humans during the Upper Paleolithic, indicating a long-standing host–virus association independent of Neanderthal transmission.

## Introduction

Human papillomaviruses (HPVs) constitute a highly diverse group of double-stranded circular DNA viruses belonging to the family *Papillomaviridae* [1]. HPVs belong in Group I of the Baltimore classification and contain a double-stranded DNA genome that relies on host cellular polymerases for replication [2]. They are non-enveloped with an icosahedral capsid enclosing a genome of approximately 8,000 base pairs, which encodes structural (L1 and L2) and non-structural (E1–E7) proteins [3,4].

Proteins L1 and L2 form the viral capsid and mediate host cell entry, while the early (E) proteins play critical roles in viral replication, episomal maintenance, and modulation of the host cell cycle [5]. HPVs have a marked tropism for squamous epithelia, infecting basal keratinocytes of the skin and mucosal surfaces, typically through microabrasions that expose the basal layer. In productive infections E1 and E2 initiate replication and transcriptional control. E4 and E5 assist in viral amplification and immune evasion and oncoproteins E6 and E7 subvert tumor suppressor pathways, particularly p53 and pRb, to promote cell proliferation and persistence of the viral genome [6].

Over two hundred HPV genotypes have been identified and are classified into several genera (Alpha, Beta, Gamma, Mu, and Nu) based on molecular and biological characteristics. Among these, *Alphapapillomavirus* species are the most clinically relevant, including both low-risk types and high-risk types implicated in malignant transformation. Within this genus, HPV16 is the most prevalent and oncogenic type, accounting for the majority of cervical cancer cases and an increasing proportion of oropharyngeal and anogenital carcinomas worldwide [7]. The HPV16 genome has a highly conserved structure although with sufficient variability to be divided into four major phylogenetic lineages (A, B, C, and D) and multiple sublineages. These variants display distinct geographic distributions and differential cancer risks, suggesting a complex and long-standing evolutionary history [8]. Understanding the origin and dispersal of these lineages has been a subject of intense debate, as it provides insights into the co-evolution of the virus and its human hosts [9,10].

Phylogenomic studies indicate that papillomaviruses have co-diverged with their primate hosts over tens of millions of years [11]. However, in humans, the specific emergence of HPV16 lineages remains an open question. The evolutionary history of HPV16 inferred from modern genomic data suggests that the HPV16A lineage was introduced into modern humans via cross-species transmission from Neanderthals or Denisovans by hybridization [10]. According to this hypothesis, HPV16B, C, and D lineages co-evolved endemically with *Homo sapiens* in Africa, whereas HPV16A originated in archaic humans and spread following the dispersal of modern humans out of Africa and their subsequent admixture events. However, this hypothesis relies on molecular clock estimates inferred from contemporary viral genomes, lacking direct evidence of viral DNA in ancient human samples to evolutionary timeline calibration.

Paleogenomics enables the recovery of ancient DNA (aDNA) from archaeological remains [12]. Novel contamination-free laboratory protocols and authentication methods allowed the identification of ancient viral genomes, such as hepatitis B virus (HBV) and Variola virus, showing that many human infections have deep temporal roots [13,14]. Despite these advances, no human papillomavirus has yet been identified in ancient *Homo sapiens* genomes, leaving open key questions on the timeline of HPV16 circulation in pre-agricultural populations [15].

In the present study, we investigated the presence of HPV sequences in aDNA data from two reference genomes: Ust’-Ishim, an Upper Paleolithic *Homo sapiens* (∼45,000 years BP) from western Siberia [16], and Ötzi the Iceman (∼5,300 years BP) from the Tyrolean Alps [17]. Ust’-Ishim represents one of the earliest known modern humans to inhabit northern Eurasia, shortly after the major distinct waves of human expansion out of Africa. Using reference-guided mapping combined with aDNA authentication criteria, we provide the first molecular evidence of HPV16 in ancient anatomically modern humans. Detection of HPV16A-related sequences in these individuals could provide a physical anchor for the viral evolutionary timeline.

## Results

### Taxonomic assessment of raw genomic data

A total of 5,751,356,775 sequencing reads were obtained, amounting to 168.9 GB of raw genomic data. A summary of this dataset, showing the contributions made by each ancient individual and sequencing platform is provided in **Table 1**.

**Table 1.** Summary of the raw sequencing datasets analyzed in this study. Data was retrieved from the NCBI SRA and ENA. Detailed run-by-run information is provided in Supplementary Table S1.

| Sample / Individual | Sequencing platform | Library layout | No. of runs | # Reads | Total data size (GB) | BioProject |
| --- | --- | --- | --- | --- | --- | --- |
| Ötzi | Illumina HiSeq 4000 | Paired-end | 5 | 1,386,308,421 | 21.9 | PRJEB5657 |
| Ötzi | AB SOLiD 4 System | Paired-end | 6 | 2,325,749,260 | 112.0 | PRJEB2830 |
| Ust'-Ishim | Illumina HiSeq 2500 | Paired-end | 1 | 2,039,299,094 | 35.0 | PRJEB6622 |
| Total | — | — | 12 | 5,751,356,775 | 168.9 | — |

The Ötzi genome contributed most of the raw reads (3.71 billion) divided across 11 runs and two platforms: five Illumina HiSeq 4000 runs (PRJEB56570) and six AB SOLiD 4 System runs (PRJEB2830). The Ust’-Ishim genome was represented by a single, high-coverage Illumina HiSeq 2500 run (ERR566093, PRJEB6622), which accounted for over 2.03 billion reads.

Detailed metadata for each of the 12 individual runs, including specific run accession numbers, sequencing platforms, and individual data volumes, is presented in **Supplementary Table 1** for comprehensive traceability.

Following data acquisition, an exploratory taxonomic assessment was conducted on each sequencing run using the SRA Run Browser’s internal classification tools. This initial step provided a tentative assignment of reads to organisms and was used to screen the raw data for the presence of reads tentatively classified within the *Papillomaviridae* family prior to formal mapping.

The results of this screening, alongside the read classification percentages, are presented in **Table 2**. The proportion of reads successfully classified (identified) varied significantly between platforms. Illumina runs exhibited higher classification rates, ranging from 57.03% to 65.30%. Conversely, the SOLiD runs showed lower classification percentages (ranging from 0.00% to 29.66%), which is consistent with the characteristics of the SOLiD platform and its shorter read lengths.

**Table 2.** Detection of papillomavirus genome fragments and overall read classification per sequencing run. Percentages indicate the proportion of reads successfully classified (identified) or remaining unclassified according to the taxonomic analysis. “Papillomavirus detected” shows whether reads assigned to the family *Papillomaviridae* were identified (“Yes”), absent (“No”), or unavailable due to missing taxonomic data (“Not available”). (*) Detailed papillomavirus types and sublineages for the positive run are displayed in the **Supplementary Figures S1–S2** (Taxonomic summaries NCBI SRA Run Browser captures) and **S3–S4** (Krona hierarchical taxonomic visualizations).

| Sample / Individual | Run accession | Sequencing platform | % reads identified | % reads unidentified | Papillomavirus detected * |
| --- | --- | --- | --- | --- | --- |
| Ötzi | ERR11837083 | Illumina | 57.81% | 42.19% | Yes |
| Ötzi | ERR11837081 | Illumina | — | — | Not available |
| Ötzi | ERR11837082 | Illumina | 57.03% | 42.97% | Yes |
| Ötzi | ERR11837084 | Illumina | 64.92% | 35.08% | Yes |
| Ötzi | ERR11837085 | Illumina | 65.30% | 34.70% | Yes |
| Ötzi | ERR107307 | SOLiD | 14.67% | 85.33% | Yes |
| Ötzi | ERR107308 | SOLiD | 15.81% | 84.19% | Yes |
| Ötzi | ERR107309 | SOLiD | 10.61% | 89.39% | Yes |
| Ötzi | ERR069107 | SOLiD | 0.00% | 100.00% | No |
| Ötzi | ERR069108 | SOLiD | 29.66% | 70.34% | No |
| Ötzi | ERR069109 | SOLiD | 0.00% | 100.00% | No |
| Ust'-Ishim | ERR566093 | Illumina | — | — | Not available |

Crucially, the exploratory screen detected the tentative presence of reads assigned to the *Papillomaviridae* family in seven of the Ötzi sequencing runs (four Illumina and three SOLiD runs). The specific Papillomavirus types and sublineages tentatively identified in these runs (detailed in **Supplementary Data 3–S24**, including taxonomic summaries (**Supplementary Data 3-9**) and Krona hierarchical visualizations (**Supplementary Data 10-S24**) were essential for the study design. These preliminary taxonomic profiles directly informed the selection of the viral reference sequences used in the subsequent alignment-based mapping techniques, ensuring a targeted and efficient virome characterization.

### Quality assessment of the reads

The quality of all 12 raw sequencing runs was systematically evaluated using FastQC to ensure data reliability and guide the subsequent preprocessing strategy. The full report webpages for all runs are provided in the Supplementary Material (**Supplementary Data 25–29** for Ötzi Illumina sequencing; **Supplementary Data 30-41** for Ötzi SOLiD sequencing; and **Supplementary Data 42 and 43** for Ust’-Ishim). The assessment revealed distinct quality profiles between the Illumina and SOLiD datasets.

The raw read quality from both the Ötzi HiSeq 4000 runs and the Ust’-Ishim HiSeq 2500 run demonstrated consistently high sequence quality, characteristic of the Illumina platform.

The five Ötzi Illumina runs (ERR11837081 to ERR11837085) showed excellent per-base quality. The median Phred score consistently exceeded 30 (indicating a 99.9% base call accuracy) for bases 1 through approximately 140. A minor, expected decline in quality was observed in the final 10 bases, where the median score dipped toward 28. Overrepresented Sequences module passed for all runs, confirming high complexity and low contamination.

The Ust’-Ishim dataset, consisting of a single paired-end run (ERR566093) quality profile mirrored the Ötzi data, with median Phred scores holding above 30 for the majority of the first 150 reads. The consistency of this quality profile across both Illumina instruments minimizes the need for aggressive quality-based trimming.

The six AB SOLiD 4 System runs for the Ötzi genome were assessed in two groups: runs designated ERR069107 to ERR069109 and runs designated ERR107307 to ERR107309. All SOLiD runs presented a distinct and significantly lower base quality compared to the Illumina data, reflecting the characteristics of color-space sequencing chemistry.

Based on the FastQC results and initial visualization, the three paired-end runs ERR107307, ERR107308, and ERR107309 were excluded from the final analysis. These runs exhibited substantially lower sequence complexity and more erratic quality score distribution than the ERR069107 to ERR069109 runs. Specifically, the group frequently failed the Per Base Sequence Quality check, suggesting pervasive instrument or systematic sequencing issues, which could have negatively impacted downstream alignment and consensus calling. This deliberate filtering step was taken to ensure that the ancient DNA comparisons relied only on the highest quality, most consistent sequencing data available for each platform. Thus, the three paired-end runs ERR069107, ERR069108, and ERR069109 were retained as they represented the best available SOLiD data.

### Genome mapping

The initial analysis involved mapping sequencing reads from three archaic hominin datasets, Ötzi (SOLiD), Ötzi (Illumina), and Ust’-Ishim (Illumina), against five major Human Papillomavirus (HPV) reference genomes (HPV16, HPV68b, HPV20, HPV18, and HPV5b), which were selected based on the results of the initial taxonomic screening.

Initial mapping attempts showed that the Ötzi Illumina sequencing data yielded significantly poor coverage and a small number of mapped reads, due to fragmentation and specific library preparation biases, consistent with previous observations on ancient DNA (aDNA). Consequently, the Ötzi Illumina dataset was excluded from further analysis, which proceeded using only the Ötzi SOLiD and Ust’-Ishim Illumina datasets.

The mapping of reads from Ötzi (SOLiD) and Ust’-Ishim (Illumina) against the five key HPV reference genomes demonstrated a highly uneven distribution of mapped reads. As shown in **Table 3**, Human Papillomavirus 16 (HPV16, NC_001526.4) consistently mapped the largest number of reads for both ancient individuals.

**Table 3.** HPV references mapped against genome reads of Ötzi and Ust’-Ishim archaic AMH using BBMap program.

| **Accession** | **# mapped reads** | **Genbank ID** | **Virus** |
| --- | --- | --- | --- |
| Ötzi |  |  |  |
| NC_001526.4 | 275,110 | GI:1047888727 | HPV16 |
| FR751039 | 256,374 | GI:315629559 | HPV68b |
| U31778 | 73,975 | GI:1020162 | HPV20 |
| NC_001357 | 33,357 | GI:9626069 | HPV18 |
| PPH5BCG | 14,868 | GI:222395 | HPV5b |
| Ust'-Ishim |  |  |  |
| NC_001526.4 | 60,379 | GI:1047888727 | HPV16 |
| FR751039 | 7,631 | GI:315629559 | HPV68b |
| U31778 | 7,306 | GI:1020162 | HPV20 |
| NC_001357 | 9,750 | GI:9626069 | HPV18 |
| PPH5BCG | 355 | GI:222395 | HPV5b |

The Ötzi SOLiD dataset, with 275,110 mapped reads, showed the highest count for HPV16, followed closely by HPV68b (256,374 reads). The Ust’-Ishim Illumina dataset showed a maximum of 60,379 reads mapped to HPV16, significantly exceeding all other viruses tested for this individual (**Table 3**).

Given the clear dominance of HPV16 in both datasets, a secondary, more focused mapping was performed against five specific reference genomes representing the four major HPV16 sublineages (A, B, C, and D) using the BBMap program. The results of this targeted analysis are presented in **Table 4**.

**Table 4.** HPV16 references of sublineages A, B, C and D mapped against genome reads of Ötzi and Ust’-Ishim archaic AMH using BBMap program.

| Accession | # mapped reads | Genbank ID | Virus | Description |
| --- | --- | --- | --- | --- |
| Ötzi |  |  |  |  |
| NC_001526.4 | 170,071 | GI:1047888727 | HPV16A | A1 European (Prototype) |
| HQ644234 | 123,377 | GI:399525615 | HPV16A | A4 European (Asian) |
| AF472508 | 55,876 | GI:27463075 | HPV16B | B1 African type 1 |
| AF472509 | 7,940 | GI:27463084 | HPV16C | C1 African type 2 |
| AF402678 | 47,167 | GI:29468132 | HPV16D | D3 Asian-American (AA2) |
| Ust'-Ishim |  |  |  |  |
| NC_001526.4 | 37,878 | GI:1047888727 | HPV16A | A1 European (Prototype) |
| HQ644234 | 53,770 | GI:399525615 | HPV16A | A4 European (Asian) |
| AF472508 | 3,558 | GI:27463075 | HPV16B | B1 African type 1 |
| AF472509 | 379 | GI:27463084 | HPV16C | C1 African type 2 |
| AF402678 | 3,779 | GI:29468132 | HPV16D | D3 Asian-American (AA2) |

For the Ötzi individual, the HPV16A sublineage references showed the highest affinity, with the A1 European prototype (NC_001526.4) and the A4 European (Asian) type (HQ644234) mapping 170,071 and 123,377 reads, respectively. Sublineages B, C, and D showed significantly lower read counts.

The Ust’-Ishim dataset also mapped preferentially to the HPV16A sublineage references. Interestingly, Ust’-Ishim showed a higher number of mapped reads against the HPV16A4 European (Asian) reference (53,770 reads) than the HPV16A1 European prototype (37,878 reads), suggesting potential genetic divergence or regional variant preference.

Mapping visualizations generated in Geneious Prime, which show read alignment and coverage across the HPV16A reference genomes, confirm these findings. Specifically, the alignment profiles for Ötzi and Ust’-Ishim indicate the highest coverage and mapping quality against HPV16A1 and HPV16A4, respectively. Both datasets exhibit a pronounced coverage peak between the E5 and L2 genes, resulting in elevated mean coverage values. Coverage in Ust’-Ishim is lower and more uneven, with multiple gaps, whereas Ötzi shows relatively uniform and high coverage. The results are illustrated by the robust read alignment of Ötzi against HPV16A1 and Ust’-Ishim against HPV16A4 (**Figure 1**), reflecting the quantitative coverage statistics (Ötzi: 97.1%; Ust’-Ishim: 30.4% of the reference covered).

**Figure 1.**
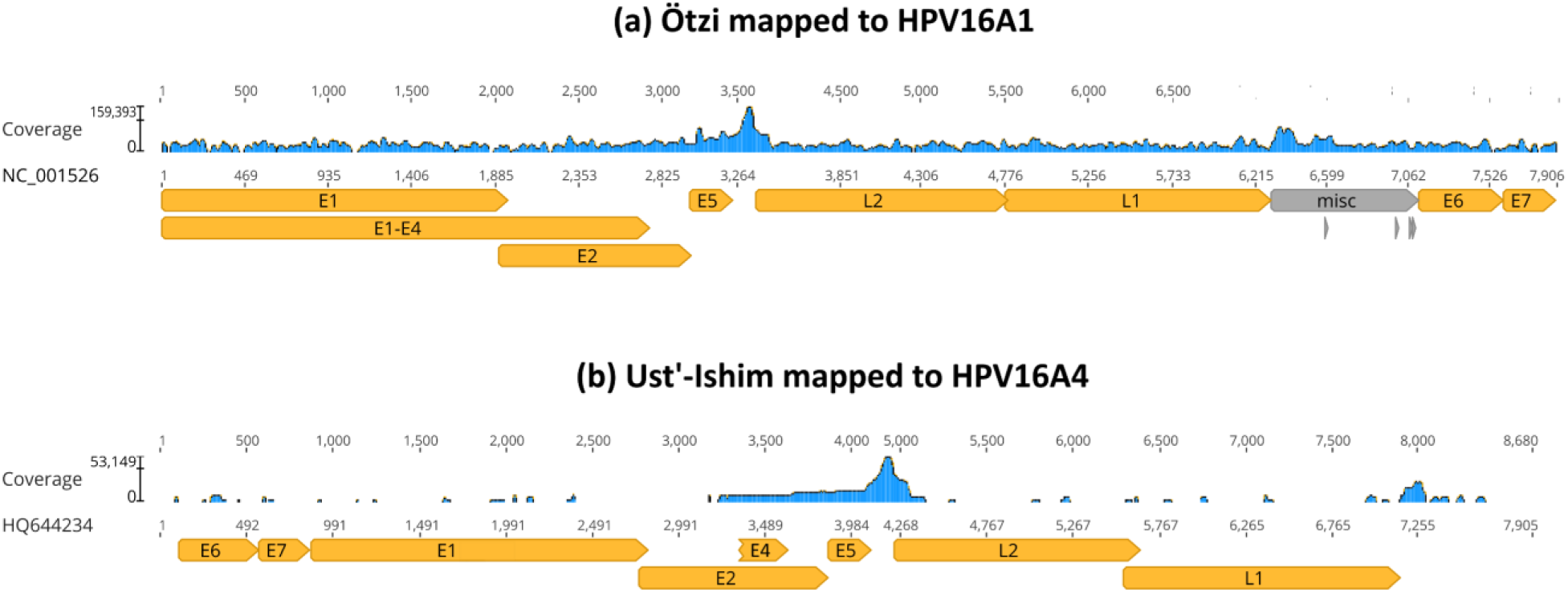
Mapping coverage of HPV16 using archaic hominin-derived reads. Reference mapping was done using BBMap. In (a) is the mapping of 170,071 Ötzi reads (mean coverage 3923.4) to the HPV16A1 reference (NC_001526.4). In (b) is the mapping of 53,770 Ust’-Ishim reads (mean coverage 2830.3) to the HPV16A4 reference (HQ644234). Orange arrows mark the protein-coding genes. Mapping was carried out simultaneously against multiple HPV16 reference sequences, as described in Materials and Methods, but only the corresponding reference is shown in each panel for clarity.

### Statistical significance of mapping

As a control for random alignment and nucleotide composition bias, mapping results against the HPV16 sublineage references were compared with those obtained using a randomized reference sequence. **Table 5** summarizes the mapping statistics (Mean Coverage, Standard Deviation, and Total Mapped Reads) for both the real sublineages and the randomized control, along with the results of the statistical analysis.

**Table 5.** Comparison between real and random reference mappings for Ötzi and Ust’-Ishim across selected HPV16 sublineages. Mean coverage, standard deviation, and number of mapped reads are reported for both real and randomized references. Welch’s *t*-tests were performed to assess the statistical significance of coverage differences, with corresponding critical values and *p*-values shown.

|  | Real reference |  |  | Random reference |  |  | Welch's *t*-test results |  |  |
| --- | --- | --- | --- | --- | --- | --- | --- | --- | --- |
|  | Coverage Mean | Coverage SD | # Reads | Coverage Mean | Coverage SD | # Reads | *t* | Critical Value | *p*-value |
| Ötzi |  |  |  |  |  |  |  |  |  |
| A1 | 3923.4 | 21728.4 | 170,071 | 23.5 | 37.1 | 5,665 | 74.0154 | 1.6449 | 0 |
| A4 | 2891.3 | 15304.6 | 123,377 | 31.8 | 87.3 | 6,706 | 65.6078 | 1.6449 | 0 |
| B1 | 890.4 | 5185.5 | 55,876 | 29.0 | 55.0 | 6,479 | 39.2479 | 1.6449 | 0 |
| C1 | 56.4 | 257.0 | 7,940 | 28.7 | 90.5 | 7,015 | 8.9937 | 1.6450 | 0 |
| D3 | 703.0 | 4625.4 | 47,167 | 24.7 | 65.9 | 5,722 | 31.8221 | 1.6449 | 0 |
| Ust'-Ishim |  |  |  |  |  |  |  |  |  |
| A1 | 1828.2 | 7252.9 | 37,878 | 0.1 | 0.3 | 11 | 49.0547 | 1.6449 | 0 |
| A4 | 2830.3 | 10711.1 | 53,770 | 0.1 | 0.6 | 23 | 61.2705 | 1.6449 | 0 |
| B1 | 92.4 | 455.0 | 3,558 | 0.0 | 0.2 | 5 | 12.1125 | 1.6453 | 0 |
| C1 | 4.0 | 26.0 | 379 | 0.1 | 0.6 | 19 | 2.9048 | 1.6488 | 0.001943 |
| D3 | 98.5 | 532.7 | 3,779 | 0.1 | 0.3 | 15 | 11.3549 | 1.6453 | 0 |

The results consistently demonstrate that the read alignments to the real HPV16 sublineage references are overwhelmingly statistically significant, confirming that the mapping success is due to specific sequence content and not random noise.

For the Ötzi sample, the difference in mapping efficiency between the real and random references was massive. The number of reads mapping to the real sublineages was orders of magnitude greater than the number mapping to the random reference. This led to extremely high *t*-values (ranging from 10.65 for C1 to 73.99 for A1) across all five comparisons. In all cases, the resulting *p*-value was 0.00 (or effectively zero), which is far below the typical significance threshold of α = 0.05. This result definitively rejects the null hypothesis that the mean coverages are equal, proving the authenticity of the Ötzi mapping.

A similar, significant pattern was observed for the Ust-Ishim sample, despite its overall lower coverage. The real reference mean coverage ranged from 4.0 (C1) to 2830.3 (A4), contrasting sharply with the consistently low mean coverage of 0.7 for the randomized reference, which only captured 118 reads.

The statistical analysis for Ust-Ishim revealed highly significant differences for sublineages A1, A4, B1, and D3, all yielding *p*-values of 0.001. Even the sublineage C1, which exhibited the lowest quantitative coverage among real references, still shows statistically significant difference with *t*-value of 2.443 and *p*-value of 0.007502, confirming it as non-random.

### Ancient DNA damage pattern

To assess the authenticity and post-mortem damage patterns of the viral reads retrieved from ancient hosts, we analyzed the BAM alignments of the individuals mapped against the HPV16 A1 and A4 reference genomes using mapDamage. The resulting plots (Stats_out_MCMC_post_pred) for Ötzi and Ust’-Ishim (**Figure 2**) revealed the characteristic patterns expected for ancient DNA, marked by an increase in cytosine deamination near read termini, producing C-to-T substitutions at the 5′ ends and complementary G-to-A substitutions at the 3′ ends.

**Figure 2.**
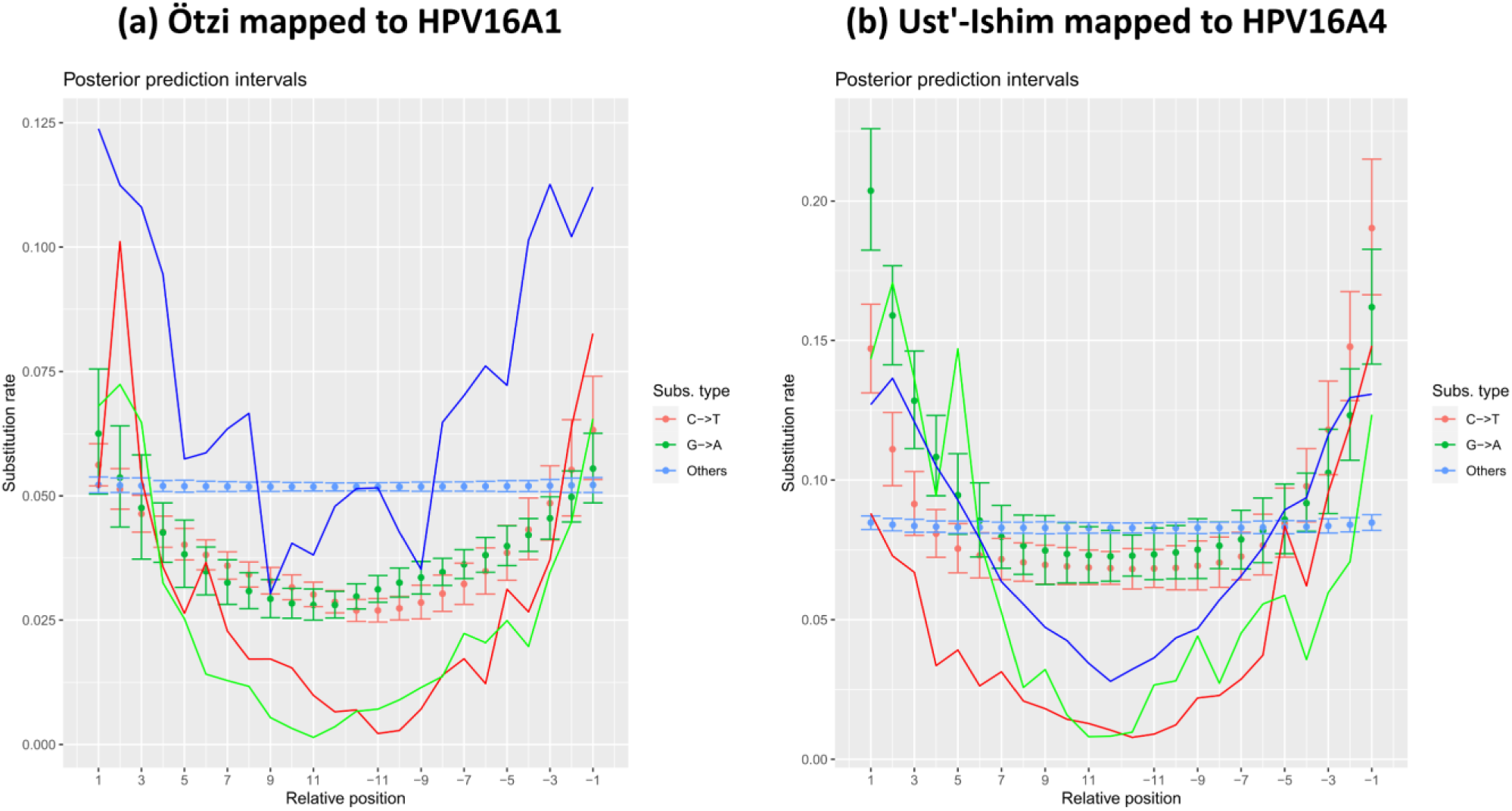
Evidence of ancient origin of reads included in the assemblies. Damage patterns were assessed with mapDamage for the Ötzi (a) and Ust’-Ishim (b) HPV16 A1 and A4 assemblies, respectively. The plots show the relative frequency of nucleotide substitutions at read termini. Colored lines represent the likelihood estimates. All assemblies exhibit the characteristic C-to-T and G-to-A substitution patterns at fragment ends, consistent with the deamination damage typically observed in ancient DNA. These terminal changes arise when cytosine deaminates to uracil (read as thymine), and guanine deaminates to xanthine, which is sequenced as adenine, using standard aDNA methods.

In both individuals, the damage plots exhibited a mild but discernible “U-shaped” pattern in the frequency of terminal misincorporations, consistent with partial cytosine deamination and fragment end damage typically observed in authentic ancient DNA molecules. The shape was less pronounced than in highly degraded specimens, possibly reflecting the low viral load, short read lengths, and variable preservation of viral fragments within host-derived libraries.

These results support that at least a subset of the mapped reads retain biochemical signatures compatible with ancient viral DNA, reinforcing the endogenous nature of the sequences rather than modern contamination.

Complementing this assessment, a specific quantitative analysis was performed using aDNA-BAMPlotter (**Figure 3**) to further validate the alignment metrics and damage profiles. For the Ötzi individual, the analysis confirmed the high integrity of the consensus assembly, showing a 93.92% genome coverage (170,071 mapped reads) accompanied by a distinct 5′ C-to-T deamination frequency of approximately 10% (0.10). Similarly, despite the lower genomic coverage retrieved for Ust’-Ishim (21.97%) and the reduced number of high-quality mapped reads, the damage signal remained robust, with a 5′ C-to-T substitution rate of approximately 8% (0.08). These quantitative metrics corroborate the mapDamage findings, statistically anchoring the ancient origin of the viral reads despite the differences in sequencing depth between the two individuals.

**Figure 3.**
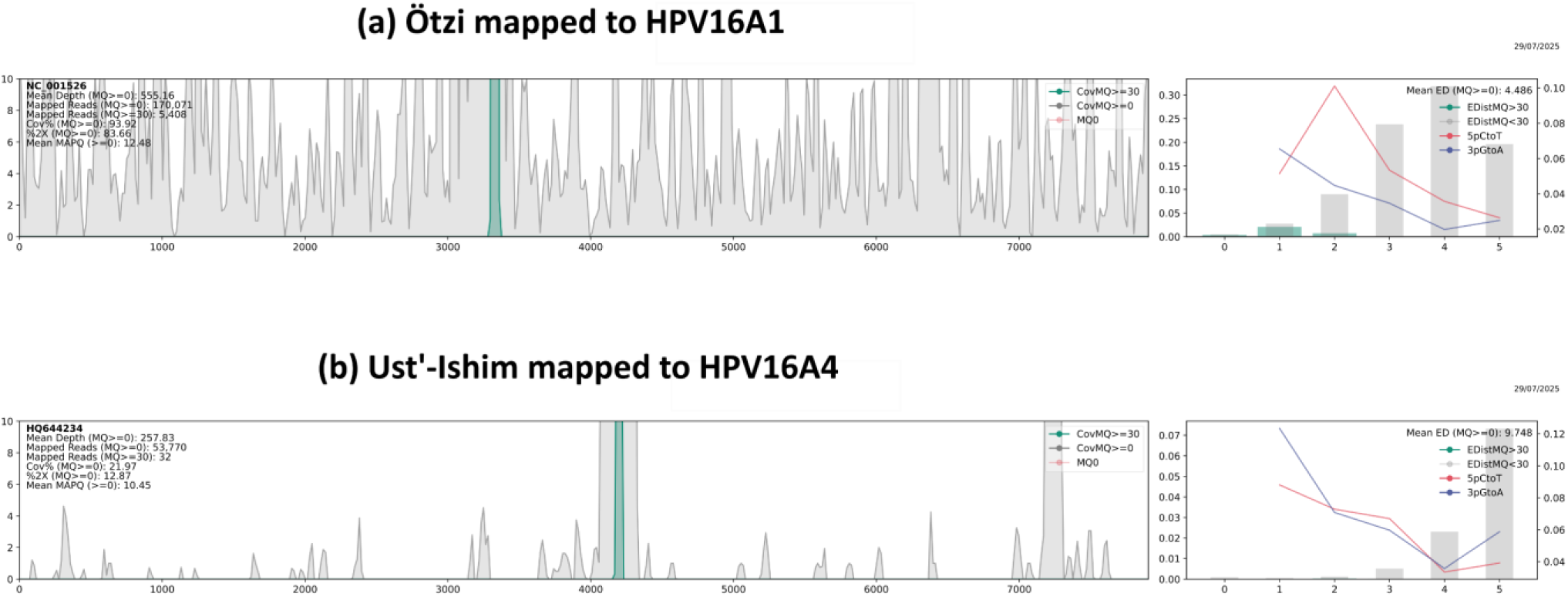
Ancient DNA damage profiles of viral reads mapped to HPV16 reference genomes. The panels display the nucleotide misincorporation patterns generated by aDNA-BAMPlotter [30] for (a) Ötzi reads aligned to the HPV16 A1 reference (NC_001526.4) and (b) Ust’-Ishim reads aligned to the HPV16 A4 reference (HQ644234). In each plot, the x-axis represents the position from the read termini (5′ end on the left, 3′ end on the right), and the y-axis indicates the frequency of nucleotide substitutions. The curves illustrate the characteristic post-mortem damage signatures: an increase in cytosine to thymine (C-to-T) transitions at the 5′ overhangs (red lines) and guanine to adenine (G-to-A) transitions at the 3′ overhangs (blue lines). The accumulation of these deamination signals at the fragment termini confirms the ancient origin of the retrieved viral sequences.

### Comparative analysis of reconstructed HPV16 sublineages

The independent multiple sequence alignments, illustrating the ancient consensus sequences alongside the HPV16 sublineage references, are presented in **Figure 4**. Based on these alignments, pairwise distance matrices were calculated to provide a quantitative assessment of the genetic identity between the ancient samples and modern viral lineages; the complete matrices are available in **Supplementary Data 44**.

**Figure 4.**
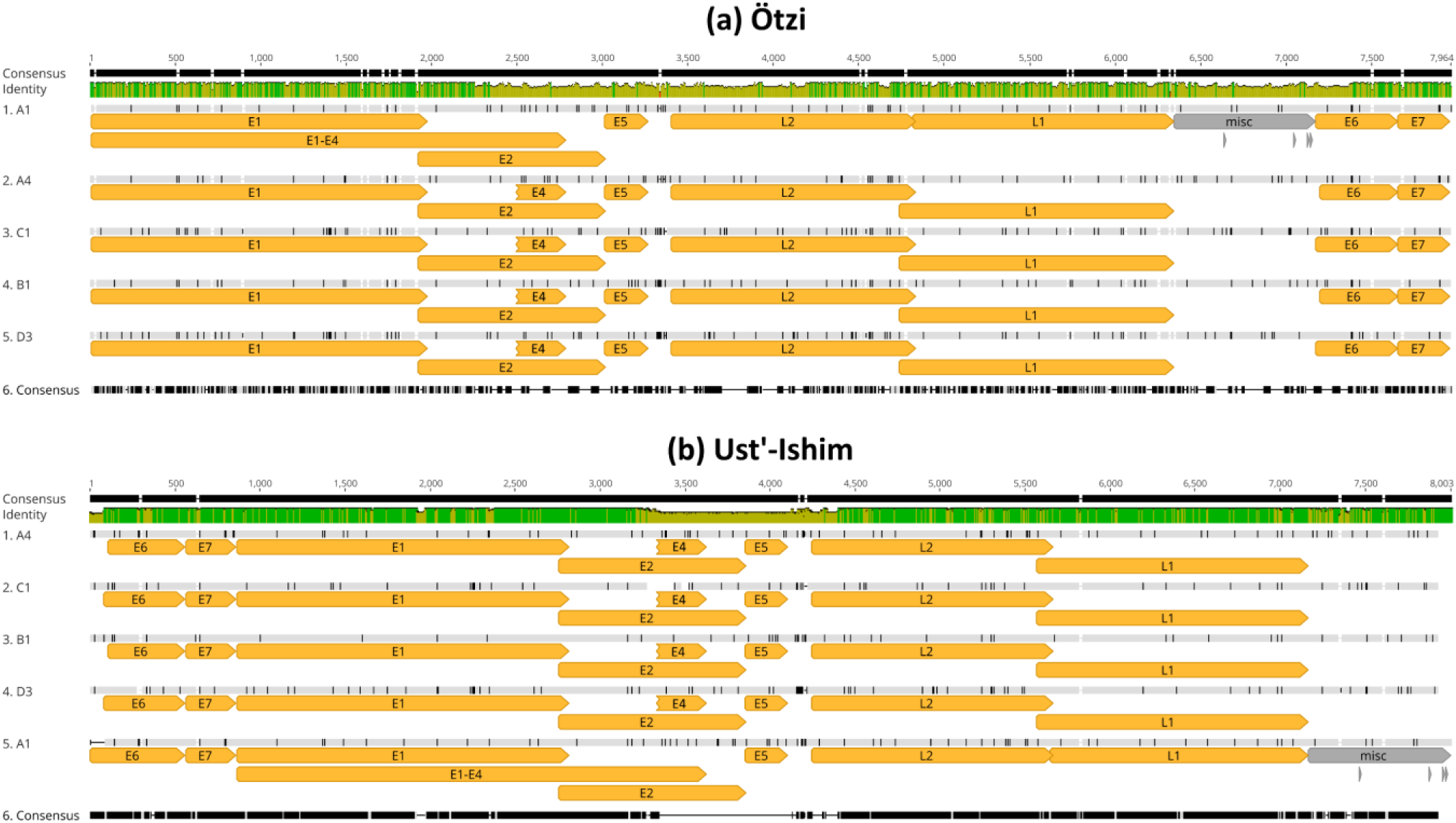
Alignment of ancient HPV16 consensus sequences with modern reference sublineages. In (a), the Ötzi HPV16 A1 consensus sequence is aligned with reference genomes representing the five HPV16 sublineages (A1, A4, B1, C1, D3). In (b), the Ust’-Ishim HPV16 A4 consensus sequence is aligned against the same set of reference sublineages. The green identity graph displayed above the tracks represents nucleotide conservation across all aligned sequences; high plateaus indicate 100% identity, while dips denote polymorphic regions. In the sequence tracks, grey shading indicates nucleotides identical to the consensus/reference, while black vertical bars highlight mismatches, representing single nucleotide polymorphisms (SNPs) or lineage-specific variations. Gaps or missing data due to low coverage are represented by dashes. Alignments were generated using MAFFT.

For the Ötzi individual, the consensus sequence generated based on the A1 mapping displayed the highest degree of genetic identity with the A1 reference genome (62.39%), followed closely by the A4 reference (61.97%). The similarity to the non-European sublineages was lower, ranging from 61.43% (B1) to 61.02% (D1). This differential identity, although influenced by the partial recovery of the viral genome, confirms the closest affinity of the Ötzi-derived virus to the A1 sublineage.

Regarding the Ust’-Ishim individual, the consensus sequence constructed using the A4 reference yielded a similar pattern of lineage-specific affinity. The Ust’-Ishim consensus shared the highest percentage of identity with the A4 sublineage reference (46.00%). Comparisons with other sublineages revealed lower identity values, with the A1 reference showing 45.34% similarity, and the African (B1, C1) and Asian-American (D1) lineages showing between 44.47% and 44.88%. Despite the lower overall genomic coverage recovered for Ust’-Ishim compared to Ötzi, the pairwise distance analysis consistently supports the classification of this viral variant within the A4 sublineage.

Across both individuals, the matrices showed the same broader structure: high similarities within the A-lineage clade and reduced identities relative to the more phylogenetically distant B, C and D references. Importantly, each ancient consensus clustered most closely with the sublineage used during read mapping, supporting the robustness of the consensus-building parameters and indicating that the resulting sequences capture the dominant viral genotype present in each host. Together, these findings confirm that the reconstructed consensus sequences from Ötzi and Ust’-Ishim faithfully reflect their respective sublineage affiliations, A1 for Ötzi and A4 for Ust’-Ishim, providing a coherent basis for subsequent comparative and evolutionary analyses. It is important to notice that pairwise identity values are reduced by missing data (‘?’) and should not be interpreted as true percent identity.

### Identification in E6 and E7 sequence variants

To investigate potential nucleotide and amino acid substitutions in the E6 and E7 oncogenes, we analyzed the variants detected in Ötzi (mapped against HPV16 A1) and Ust’-Ishim (mapped against HPV16 A4). Given the fragmented nature of the ancient DNA, variant calling was performed with relaxed parameters to maximize sensitivity. To distinguish likely biological variants from post-mortem damage and sequencing noise, we prioritized non-synonymous mutations that displayed higher variant frequencies or coverage support. The complete dataset of all identified non-synonymous changes is provided in **Supplementary Data 45**.

A subset of the most relevant (coverage ⋝ 5x; variant frequency ⋝ 15%) non-synonymous amino acid substitutions found in Ötzi is summarized in **Figure 5** (for full figure including all non-synonymous variants see Supplementary Figures S1 (E6) and S2 (E7)). In the E6 gene, we observed several missense mutations with variant frequencies ranging from 16.7% to 42.9%. Notably, position 7218 showed a D-to-H (Aspartic acid to Histidine) substitution with 28.6% frequency supported by 7 reads, a transversion less likely to be caused by deamination damage. High-frequency transitions were also observed at positions 7371 (Y-to-H, 40%) and 7387 (Y-to-C, 42.9%).

**Figure 5.**
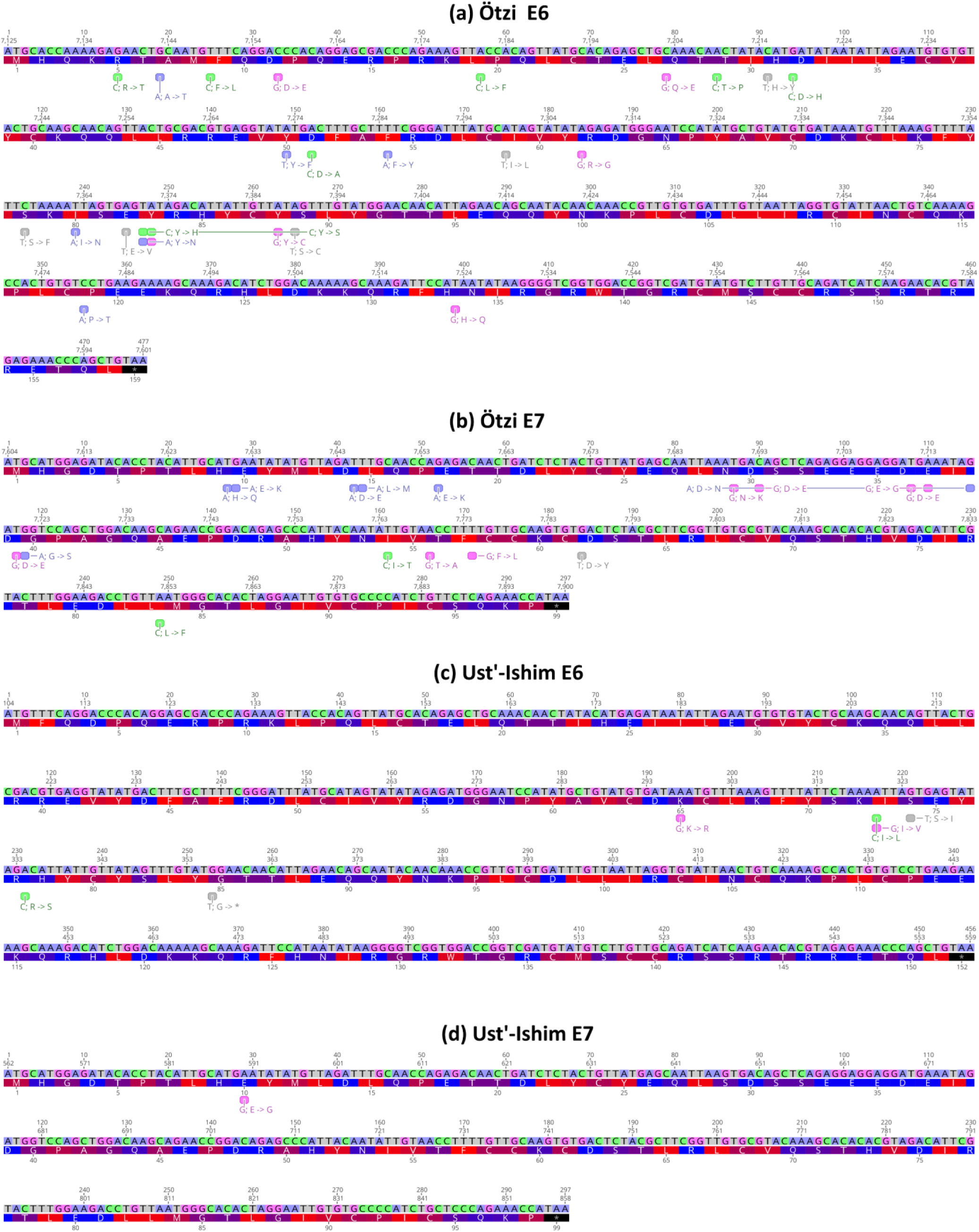
Non-synonymous SNPs identified in the HPV16 E6 and E7 genes of the ancient genomes compared against their respective sublineage reference sequences. The panels display the variations found in: (a) Ötzi E6 and (b) Ötzi E7, compared against the A1 reference (NC_001526.4); and (c) Ust’-Ishim E6 and (d) Ust’-Ishim E7, compared against the A4 reference (HQ644234). Nucleotide colors follow the same scheme: A=purple, C=green, G=pink, and T=gray. Amino acid residues are colored according to hydrophobicity, with red representing the most hydrophobic residues (value=0), blue representing the most hydrophilic (value=0.5), and purple indicating intermediate hydrophobicity (value=1), following the Expasy scale (https://web.expasy.org/protscale/pscale/Hphob.Black.html). For each site, the reference nucleotide is shown, and the altered base observed in the ancient sequence is highlighted by a colored box. The resulting amino-acid substitution is annotated directly beside the modified position.

The analysis of the E7 gene in Ötzi yielded variants supported by higher read depths compared to E6. Several transversions were identified with robust coverage, reinforcing the authenticity of the signal. Significant variants included substitutions at positions 7690 (N-to-K, 21.1% frequency, 19x coverage), 7711 (D-to-E, 20.8% frequency, 24x coverage), and 7720 (D-to-E, 22.2% frequency, 9x coverage). The detection of these transversions at high coverage suggests biological variation rather than post-mortem artifacts. Additionally, position 7852 showed an L-to-F substitution with 28.6% frequency.

The Ust’-Ishim dataset exhibited significantly lower coverage across the viral genome compared to Ötzi, resulting in a sparse yield of variants (**Figure 5**). Despite the low depth, distinct high-frequency variants were observed in the E6 oncogene. Notably, position 324 presented an S-to-I (Serine to Isoleucine) substitution with 60% frequency supported by 5 reads. Position 320 showed a complex polymorphism where the reference Isoleucine (I) was substituted by Leucine (L) or Valine (V) in 40% of the reads (combined frequency), also supported by 5 reads. Other potential variants, such as K-to-R at position 297 (E6) and E-to-G at position 590 (E7), were identified with 100% and 50% frequencies respectively, though supported by extremely low coverage (<3 reads), warranting caution in their interpretation.

Although the variable depth of coverage prevents a definitive determination of fixed mutations in these ancient viral populations, the presence of these non-synonymous variants suggests sequence divergence from their respective modern references. These observed substitutions in the E6 and E7 oncogenes were evaluated for their potential impact on protein function and oncogenicity.

## Discussion

The present study provides a multi-layered analytical framework for detecting, validating, and characterizing ancient HPV16 sequences recovered from two well-known ancient individuals, Ötzi and Ust’-Ishim. By integrating taxonomic screening, targeted viral mapping, statistical validation using randomized controls, ancient DNA damage profiling, and comparative genomic analyses, we demonstrate that both individuals carried HPV16 variants belonging to the A lineage, with Ötzi affiliating most closely with sublineage A1 and Ust’-Ishim with sublineage A4. The convergence of evidence across these independent methodological approaches strongly supports the authenticity and biological relevance of the recovered viral sequences, despite the inherent challenges of ancient viral DNA recovery.

The initial mapping step revealed a clear predominance of HPV16 over other tested HPV types in both individuals, indicating that the detected signal reflects specific viral presence rather than spurious alignment. The exclusion of the Ötzi Illumina dataset due to insufficient viral mapping further ensured that subsequent analyses were based on the most informative and reliable data. Mapping against sublineage-specific HPV16 references refined this observation and produced consistent patterns: the Ötzi SOLiD reads mapped preferentially to HPV16 A1, whereas Ust’-Ishim Illumina reads aligned most strongly to HPV16 A4. These affinities are biologically coherent in light of known HPV16 phylogeography, where A1 predominates in European populations and A4 is associated with early Eurasian groups [9,10].

The predominance of HPV16 in both ancient individuals is itself an informative finding. Among all high-risk HPV genotypes, HPV16 is consistently the most prevalent worldwide and exhibits the highest oncogenic potential. Its detection in two genetically and geographically distinct Late Pleistocene and Holocene individuals suggests that HPV16 has maintained a long-standing association with human populations and may have undergone host-specific evolutionary stabilization long before the emergence of modern epidemiological patterns. The recovery of HPV16, rather than other high-risk types such as HPV18, HPV31, or HPV45, aligns with the genotype’s known global ubiquity and supports the hypothesis that HPV16 has been the dominant circulating high-risk lineage across diverse human groups for millennia [8–10].

Our previous analysis of Neanderthal sequencing data demonstrated that the papillomavirus recovered from Chagyrskaya Neanderthals belongs to HPV12, not HPV16 [18]. This suggests that Neanderthals were not a reservoir of HPV16 and further weakens the hypothesis that the HPV16A-related lineage detected in Ust’-Ishim and Ötzi originated via viral introgression from archaic hominins. Instead, the detection of HPV16A-related sequences in both individuals suggests infections circulating within anatomically modern humans. These findings indicate that HPV16 was already present in *Homo sapiens* and support a scenario in which the deep history of HPV16 lineage A predates the contact with Neanderthals.

To ensure that the observed mapping signals could not be attributed to random alignment or nucleotide composition bias, a particular concern in analyses involving short, damaged ancient DNA fragments, we compared the mapping metrics obtained from real references to those obtained from randomized control sequences. The stark differences in mean coverage, mapping depth, and total mapped reads resulted in extremely high t-values and p-values effectively equal to zero across all comparisons. These results decisively reject the hypothesis that the viral alignments could be explained by stochastic noise. Notably, the statistical separation remained significant even in the lower-coverage Ust’-Ishim dataset, indicating that the biological signal persists despite greater fragmentation and uneven representation of viral genomic regions.

An independent line of evidence supporting authenticity comes from the detection of characteristic ancient DNA damage patterns. The mapDamage analysis revealed a mild but consistent enrichment of C-to-T substitutions at 5′ termini and G-to-A substitutions at 3′ termini, forming the expected “U-shaped” misincorporation profile associated with cytosine deamination. Although less pronounced than in highly degraded ancient human genomic libraries, this pattern aligns with expectations for low-abundance viral DNA embedded within larger host-derived sequencing datasets. The detection of such damage in both individuals, despite differences in preservation and sequencing platform, reinforces the endogenous nature of the viral fragments and argues against modern contamination in relation to HPV16 in the sequences.

Although exogenous contamination was detected in the SOLiD dataset of the Ötzi individual, multiple lines of evidence indicate that this contamination did not extend to the HPV16 reads. First, mapDamage profiles generated from the HPV16-aligned reads show the canonical signatures of ancient DNA, cytosine deamination at 5’ ends and complementary G-to-A misincorporations at 3’ ends, consistent with authentic ancient molecules rather than modern contaminants. Second, the reads contributing to the HPV16 mapping displayed fragment-length distributions in the expected aDNA range, with no enrichment of longer, undamaged fragments that would be indicative of modern viral DNA. Third, the consensus sequence exhibited internally coherent positional damage and no clustering of low-complexity or anomalous reads that might suggest contamination-driven artefacts. Together, these indicators confirm that, although the SOLiD run contains contaminants at the global dataset level, the HPV16-positive fraction is composed of endogenous, ancient molecules and is not attributable to modern or laboratory-derived viral DNA.

Furthermore, the quantitative metrics derived from the aDNA-BAMPlotter analysis substantiate the integrity of these assemblies. For the Ötzi sublineage, the extensive genomic coverage (93.92%) combined with a distinct ∼10% deamination frequency establishes this sequence as a high-confidence, near-complete ancient viral genome. Conversely, in the more fragmented Ust’-Ishim dataset, the persistence of a ∼8% terminal deamination signal provides a critical validation filter. It confirms that despite the lower coverage (21.97%) and mapping quality scores, the aligned reads represent genuine ancient viral templates rather than modern environmental background or sequencing artifacts.

Comparative genomic analyses further support these interpretations. Pairwise identity matrices constructed from the recovered consensus sequences show that each ancient HPV16 genome is most like the sublineage that produced the highest mapping scores. Ötzi’s consensus aligns most closely with the A1 reference, while that of Ust’-Ishim aligns most strongly with A4, despite lower coverage and greater sequence incompleteness. Importantly, both consensus sequences display the same hierarchical pattern: higher similarity to the A lineage and progressively lower similarity to the more distantly related B, C, and D lineages. This internal consistency across coverage levels, mapping statistics, and evolutionary distances indicates that the consensus sequences reliably capture the dominant viral genotype present in each host.

A potential challenge in ancient viral genomics is reference bias, where the choice of a modern template influences the resulting consensus sequence. However, our methodology mitigated this by employing a competitive mapping strategy prior to consensus generation. As observed in the mapping results, the Ust’-Ishim dataset exhibited a clear preferential affinity for the A4 sublineage (53,770 reads) over the A1 prototype (37,878 reads), despite the high sequence similarity between these European variants. This differential read recruitment indicates that the sublineage classification was driven by endogenous biological variation present in the ancient reads rather than an artifact of the assembly process. Consequently, the resulting pairwise distances, which confirmed the A1 status of Ötzi and A4 status of Ust’-Ishim, reflect genuine phylogenetic signals.

Taken together, these findings support a coherent picture in which the reconstructed viral sequences represent authentic ancient HPV16 genomes that reflect the long-term association between papillomaviruses and human populations. The presence of lineage A1 in Ötzi is consistent with its current predominance in European populations, while the identification of lineage A4 in Ust’-Ishim is compatible with early dispersals of anatomically modern humans into Eurasia. This study thus expands the evolutionary and temporal context of HPV16, demonstrating its presence in ancient Eurasian individuals separated by more than 5,000 years and highlighting the potential for reconstructing ancient viral genomes even from low-level, partially preserved DNA fragments.

The phylogenetic placement of these ancient genomes aligns with current models of HPV16 evolution. The identification of an HPV16 A1 genome in Ötzi (3,300 BCE) provides direct molecular evidence that the A1 sublineage was already established in Central Europe during the Copper Age. Conversely, the classification of the Ust’-Ishim virus (45,000 BP) as A4, a sublineage often associated with Asian-European dispersal, supports the hypothesis of an ancient divergence between the A1 and A4 clades. The recovery of an A4-like sequence from a Paleolithic Siberian individual suggests that the ancestral diversification of the A lineage predates the Neolithic transition, potentially accompanying early human migrations across Eurasia.

In typical ancient pathogen workflows, duplicate removal is not recommended for highly degraded viral datasets, particularly when working with single-stranded or very short double-stranded fragments [12,19]. Because viral genomes are small, true biological duplicates are common and expected, and aggressive duplicate filtering can disproportionately eliminate authentic molecules, artificially reduce coverage, and bias downstream estimates of variant frequency. For this reason, consensus generation and sublineage classification in ancient virology rely on retaining all reads, including duplicates, while instead applying stringent mapping, quality, and damage-based authenticity filters.

Nevertheless, to evaluate the robustness of our results, we performed an auxiliary duplicate-removal pipeline for both individuals, documented in **Supplementary Data 46**. Despite the expected decrease in coverage, pairwise distance patterns, variant calls, and sublineage affinity remained unchanged. A Welch’s *t*-test comparing identity percentages from the original and deduplicated datasets showed no statistically significant difference, confirming that the inclusion of duplicates did not inflate or distort the conclusions. This further supports the stability of the reconstructed consensus sequences and their correct assignment to the A1 (Ötzi) and A4 (Ust’-Ishim) sublineages.

The analysis of the E6 and E7 coding regions in the HPV16 genomes recovered from Ötzi and Ust-Ishim revealed considerable apparent variability, but most putative substitutions are best interpreted within the constraints imposed by ancient DNA damage and extremely low coverage at many sites. In Ötzi, the PDZ-binding motif of E6 (ETQL) [20], a functionally critical C-terminal element required for PDZ-domain protein degradation, was covered by a single read and contained a synonymous third-position transversion (CTG-to-CTT). Because the substitution does not alter the encoded amino acid and falls at a position highly susceptible to ancient DNA misincorporation, there is no evidence that the PDZ-binding function was altered in the viral sequence infecting Ötzi. In Ust-Ishim, however, this C-terminal region had no coverage, preventing any assessment of substitutions within the ETQL motif. More broadly, Ust-Ishim exhibited consistently low or absent coverage across large portions of E6 and E7, a limitation that recurs throughout the gene and constrains the interpretation of multiple regions discussed below. As a result, sequence preservation can only be evaluated for positions that are covered in each individual; for Ust-Ishim, many sites remain undetermined rather than demonstrably conserved.

The high-affinity p53-binding region of E6 (residues 113–122) [21], essential for the degradation of p53 by the E6–E6AP complex, also showed numerous putative mutations in Ötzi, many appearing at very low frequencies and across positions with coverage values at or near the detection threshold. Several positions displayed mutually exclusive codon changes (e.g., both A-to-G and A-to-C producing incompatible CAG variants at the same site), and occasional truncation calls occurred alongside amino acid substitutions. This pattern is characteristic of ancient DNA damage, low-/complexity or repetitive sequence effects, and misaligned reads rather than genuine biological heterogeneity. In Ust-Ishim, coverage across this region was limited, but in the few positions with adequate data, no supported nonsynonymous substitutions were observed. Because most positions lacked sufficient depth to evaluate, the absence of substitution cannot be taken as evidence of true conservation in Ust-Ishim; rather, the available reads simply did not show credible variation. Accordingly, while there is no positive evidence of disruption to the p53-binding region in either genome, conservation can only be confidently assessed in the limited positions with reliable coverage. The zinc-finger domains of E6, which stabilize E6 structure and mediate multiple oncogenic interactions [22], displayed some of the most irregular patterns in Ötzi. Positions C37, C40, C70, and C73 showed multiple low-frequency substitutions—including Cys-to-Ser/Trp and Cys-to-Phe changes, which would be structurally disruptive if real. However, these calls consistently arose at depths of 1–2× and often produced contradictory codons at the same position (e.g., CCT-to-ACT, CCT-to-GCT, CCT-to-CAT each at ∼10–20%), an inconsistent pattern incompatible with true quasispecies variation in a small DNA virus with low intrahost diversity. Ust-Ishim provided partial coverage across this region; the zinc-finger cysteines that were covered appeared without supported amino acid substitutions. Nevertheless, several cysteine positions lacked data entirely, meaning that true conservation cannot be inferred genome-wide for Ust-Ishim. Notably, the deeply constrained residue C110 was covered in both genomes and showed no mutation, supporting the maintenance of this critical structural site.

The linker region between the two E6 zinc fingers displayed similar patterns in Ötzi, with mutually exclusive codon calls and low-frequency truncations across positions with minimal depth. Although this region is less strictly constrained than the zinc-coordinating residues, the absence of coherent, high-confidence variants—and the lack of convincing substitutions in the few positions covered in Ust-Ishim—suggest that the ancestral HPV16 A lineage infecting these individuals likely retained a typical E6 architecture. However, as with other regions, positions without coverage in Ust-Ishim cannot be assumed to be conserved.

In E7, both ancient genomes recovered at least partial coverage of the canonical LxCxE motif (residues 22–26) [23,24], the essential domain for pRB binding and disruption of cell-cycle control. Ötzi exhibited two variant calls near this motif, including a single-read truncation and an E-to-Q change, but both occurred at extremely low depth and displayed patterns consistent with sequencing noise. In Ust-Ishim, coverage across this region was limited but did not show credible amino acid changes in the reads available; however, lack of coverage at several sites prevents asserting full motif conservation. Across CR1–CR3, Ötzi again showed scattered low-frequency substitutions typical of ancient DNA artefacts, while Ust-Ishim’s sparse but cleaner data did not reveal supported nonsynonymous changes at the positions it covered. The CKII phosphorylation sites (S31/S32), which modulate E7 stability and oncogenic potency, were also affected by extremely low depth in Ötzi, generating several low-frequency substitutions (S-to-N, S-to-R, S-to-P) that are unlikely to represent real biological changes; in Ust-Ishim, these sites were again poorly covered and therefore cannot be evaluated.

Taken together, the comparative analysis of Ötzi and Ust-Ishim indicates that despite numerous apparent substitutions in Ötzi, most matching expected signatures of ancient DNA damage, the limited coverage in both genomes, and especially the extensive missing data in Ust-Ishim, constrains definitive conclusions about the full conservation of functional motifs. Where coverage is sufficient (e.g., C110 in E6, portions of the zinc-finger cores, and segments of E7’s CR1–CR3), neither genome shows high-confidence deviations from canonical HPV16 A-lineage sequences. However, wide regions of E6 and E7 in Ust-Ishim are effectively indeterminate, meaning the absence of mutations should not be interpreted as evidence of preservation. Overall, the combined data suggests that there is no credible evidence for naturally occurring attenuating or hyper-oncogenic mutations in the positions that could be robustly evaluated, but substantial portions of both genes, particularly in Ust-Ishim, remain unresolved due to insufficient depth. This underscores both the power and the current limitations of ancient viral genomics, while reinforcing the value of conservative, damage-aware interpretation frameworks for reconstructing the deep evolutionary history of oncogenic viruses.

In conclusion, this study presents direct molecular evidence of Human Papillomavirus type 16 (HPV16) in anatomically modern humans, recovering viral sequences from the genomes of Ust’-Ishim (∼45,000 BP) and Ötzi the Iceman (∼5,300 BP). Using a workflow involving reference-guided mapping and ancient DNA authentication, we observed that sequences related to oncogenic HPV lineages were preserved in these Eurasian samples. Our findings offer a physical calibration point for the evolutionary timeline of *Papillomaviridae*. The identification of HPV16, specifically affinities with the A-lineage, in the Ust’-Ishim individual adds a new layer of complexity to the understanding of viral dispersal. These data are consistent with the possibility that the presence of HPV16A in *Homo sapiens* could predate the Neolithic transition, potentially indicating a host–virus association established earlier than previously inferred from molecular clock estimates alone. Consequently, these results invite a reassessment of current theoretical models, hinting that the dynamics of archaic-to-modern transmission or the virus’s ancestral geography might be more intricate than the prevailing introgression hypothesis suggests. The recovery of viral reads from sequencing data not originally targeted for viral capture underscores the potential of archived paleogenomic datasets to illuminate the deep history of human pathogens. Finally, while the presence of viral DNA has been detected, its biological state within the host requires further investigation. Given the oncogenic potential of HPV16, which is often linked to the physical state of the viral genome, a viral integration analysis is currently underway. Future work will aim to assess whether the viral sequences existed as episomes or were integrated into the host chromatin, providing a deeper perspective on the potential pathogenicity of these ancient viral strains.

## Materials and Methods

This section describes the procedures carried out for the acquisition, processing, and analysis of the data used in this study. The objective was to establish a standardized and reproducible workflow suitable for ancient DNA research, following the methodological pipelines established in our research group and previously applied in paleovirology [18,25].

The analyses were structured into seven main stages: (i) acquisition of sequencing data from public repositories; (ii) preprocessing and quality assessment of sequencing reads; (iii) exploratory mapping against a panel of *Human papillomavirus* (HPV) reference genomes, which revealed a predominance of reads aligning to HPV16. This observation guided subsequent analyses focused on HPV16, using representative reference genomes of its major subtypes and sublineages (A, B, C, and D) detected in the ancient samples; (iv) statistical validation of the mapping results to verify that the alignment patterns were not due to random occurrence; (v) investigation of potential contamination and the assessment of characteristic ancient DNA damage patterns; (vi) multiple sequence alignment and pairwise genetic distance analyses among the identified HPV16 sequences and modern reference lineages, providing comparative insights into viral diversity and evolutionary relationships; and (vii) identification and interpretation of non-synonymous single-nucleotide variants in the E6 and E7 oncoproteins in order to evaluate their potential functional and oncogenic implications.

The selection of databases, software, and analytical parameters was based on criteria of broad scientific use, relevance to ancient genome studies, and the availability of tools enabling robust comparative analyses. Accordingly, all procedures were designed to align with best practices recommended in the literature for the processing of ancient genomic data.

Use of AI Tools: To support the interpretation of results, assist in drafting sections of the manuscript, and help identify potential errors or inconsistencies, we made use of AI language models, specifically ChatGPT (https://chatgpt.com/) and Google Gemini (https://gemini.google.com/app), during the period of February 2025 to December 2025. These tools were employed solely as writing and analytical aids, and all information generated through them was independently verified by us. All final interpretations, decisions, and conclusions remain our own.

### Data Acquisition

Illumina sequencing reads of the Ötzi genome were obtained from the European Nucleotide Archive (accessed on 6 March 2025) (https://www.ebi.ac.uk/ena/browser/home) under project accession number PRJEB56570 [26]. Five Illumina paired-end sequencing runs (ERR11837081–ERR11837085) were acquired, all publicly available in FASTQ format.

SOLiD sequencing reads of the Ötzi genome were also retrieved from the European Nucleotide Archive (accessed on 12 March 2025) under accession number PRJEB2830 [14], corresponding to six paired-end runs (ERR069107–ERR069109 and ERR107307–ERR107309).

Illumina paired-end sequencing reads of the Ust’-Ishim genome were obtained from the Galaxy platform (accessed on 24 March 2025) [27] using accession number ERR566093 [16]. The dataset was retrieved through the tool Get Data -to- Download and Extract Reads in FASTQ format from NCBI SRA (Galaxy Version 3.1.1+galaxy1), which internally reconstructs FASTQ files through the SRA toolkit. This conversion step does not guarantee preservation of the original read order and frequently produces forward and reverse FASTQ files in which several reads don’t have a corresponding pair. As a result, the two files were imported into the workflow as separate, unpaired datasets, since attempting to enforce pairing would have caused substantial data loss. This does not affect the aims of the present study, as all downstream analyses rely exclusively on single-read alignment, including quality control, reference-based mapping, variant inspection, and ancient-DNA damage assessment, making paired-end information unnecessary for the analytical framework employed.

The viral reference sequences used in this study were retrieved from the NCBI database (https://www.ncbi.nlm.nih.gov/ accessed on 2 May 2025) in GenBank format (**Tables 3 and 4**). All datasets were imported into the Geneious Prime v2025.1 software (https://www.geneious.com) for initial inspection and data organization and subsequent analysis, Ötzi’s by pairing forward and reverse files and Ust’-Ishim’s kept separate.

### Preprocessing

The quality of all sequencing reads was evaluated using FastQC (https://www.bioinformatics.babraham.ac.uk/projects/fastqc/, accessed on 29 April 2025). (Galaxy Version 0.74+galaxy1) [21] through the Galaxy platform [19]. On Galaxy, FASTQ files of each run were obtained via Get Data -to- Download and Extract Reads in FASTQ format from NCBI SRA (Galaxy Version 3.1.1+galaxy1) and from the history were used for the individual analysis using FastQC to generate read quality reports, including per-base sequence quality, GC content, and sequence duplication levels.

Each sequencing run was also individually examined in the SRA Run Browser (https://www.ncbi.nlm.nih.gov/Traces/index.html?view=run_browser&display=metadata, accessed on 2 May 2025) to confirm general sequencing information, including the total number of spots and bases, file size, sequencing platform and strategy, library type and layout, quality metrics based on Phred scores, and, when available, associated taxonomic analyses. This exploratory step was performed to guide subsequent stages of the study and to preliminarily evaluate the presence of reads assigned to the *Papillomaviridae* family. Within the Run Browser, this identification is based on automatic taxonomic classification performed by NCBI’s internal alignment tools, which provide tentative organismal assignments for reads in each sample [28]. Accordingly, this inspection offered a preliminary overview of potential viral agents without yet representing a formal mapping analysis.

### Mapping against viral reference genomes

Based on the detection of *Papillomaviridae* reads during the taxonomic screening conducted in the preprocessing stage, complete genomes of the identified papillomaviruses were retrieved from the NCBI database, prioritizing reference sequences (RefSeqs) when available. The initially selected reference genomes included *Human papillomavirus* types 16, 18, 68b, 20, and 5b. Their respective NCBI accession numbers are provided in **Table 3**.

The five *Human papillomavirus* reference genomes were simultaneously aligned against each set of sample reads, resulting in four independent mapping analyses: (i) references against Ötzi Illumina reads, (ii) references against Ötzi SOLiD paired-end reads, (iii) references against Ötzi SOLiD reads, and (iv) references against Ust’-Ishim Illumina unpaired reads.

All mappings were performed in Geneious Prime using BBMap aligner for DNA/RNAseq version 38.84 [29] as the alignment algorithm. BBMap was selected for its high tolerance to sequence length variation and typical ancient DNA damage, as well as its efficient performance in handling short and fragmented reads. All the mappings were performed with normal sensitivity, no trimming before alignment, and multiple best matches mapped randomly. Additional parameters included a k-mer length of 13 and a maximum indel size of 16,000.

Based on the results obtained, subsequent analyses were carried out using the Ötzi SOLiD paired-end and Ust’-Ishim Illumina unpaired datasets. These were remapped against sublineages of the main *Human papillomaviru*s genotype identified in the initial mappings: HPV16.

To account for the intraspecific genetic diversity of HPV16 and to ensure representation of its four major lineages (A–D), the sublineages A1, A4, B1, C1, and D3 were selected, all of which have complete genome sequences publicly available in NCBI. The corresponding accession numbers were obtained from Pimenoff *et al.* [10], who analyzed the evolutionary relationships between archaic and modern human HPV16 lineages. The A1 sublineage corresponded to the same reference used for HPV16 in the previous mapping analysis.

The subdivision of HPV16 into distinct lineages and sublineages is based on genomic divergence thresholds established by Burk *et al.* [8], who proposed that variants differing by more than 1–2% across the complete genome should be classified as separate lineages. This framework supports the inclusion of the selected sublineages to capture the known phylogenetic structure of HPV16. The reference sequences and their corresponding NCBI accession numbers are listed in **Table 4**.

All mappings were performed on a workstation equipped with a 12th Gen Intel® Core™ i9-12900K processor (24 CPUs, ∼3.2 GHz) and 128 GB of RAM.

### Statistical Analysis of Randomness

To determine if mapping of aDNA reads to HPV16 sublineage references (A1, A4, B1, C1, and D3) represent biological signal rather than by random noise (spurious sequence similarity or nucleotide composition bias), statistical negative control experiments were performed as previously described [18,25]. Using the same HPV16 length and nucleotide composition (A, C, G, and T frequencies) random reference sequences were generated with the Random Sequence Generator (UGENE) [26]. The artificial reference sequences maintained identical base composition, are randomized versions of viral references and used as negative controls in independent mappings against Ötzi (SOLiD) and Ust’-Ishim (Illumina) aDNA reads datasets.

Mapping statistics obtained with random references (number of aligned reads, mean, and standard deviation) were compared to those from the original mapping against the real HPV16 sublineage references using Welch’s *t*-test. This assesses whether two groups with unequal variances and sample sizes differ significantly in their means (https://www.statskingdom.com/t_test.html#two_t_w, accessed on 2 May 2025). The test was conducted as right-tailed, with a standardized effect size of 0.5 (medium effect), a significance level of 0.05, mean difference set to 0, and outliers retained in the analysis.

This procedure allowed us to determine whether the degree of alignment observed with the real reference genomes differed significantly from what would be expected chance alone, given a sequence with identical overall nucleotide composition. Thus, results showing significant differences (*p* < 0.05) between the real and random mappings indicate genuine genomic affinity, suggesting that the observed alignments reflect true biological correspondence between reads and HPV16 viral sequences rather than compositional similarity alone. Conversely, non-significant results suggest that the mapping could be explained by base composition rather than true homology.

### Contamination and Ancient DNA Damage Analysis

The characteristic damage patterns of ancient DNA were assessed using mapDamage v2.2.2 [28], run in a Docker container via "docker run -it -v "/path/to/Mapdamage_Data:/data" quay.io/biocontainers/mapdamage2:2.2.2--pyr43hdfd78af_0 mapDamage -i /data/otzi.bam -r/data/av.fasta -d /data" and applied to the BAM files generated from the mappings. The software estimates the frequency and positional distribution of nucleotide misincorporations associated with postmortem degradation, particularly C-to-T transitions at the 5′ ends and G-to-A transitions at the 3′ ends of DNA fragments. These patterns, widely documented in ancient DNA studies [29], represent key molecular authenticity indicators and aid in distinguishing endogenous material from potential modern contamination. The aDNA-BAMPlotter v2.0.1 [30] was used for graphical visualization and documentation of the mapped genomic regions, allowing detailed inspection of read coverage and density, as well as identification of atypical alignment patterns that could indicate contamination or technical artifacts.

### Consensus Sequence Generation, Multiple Sequence Alignment, and Distance Estimation

Consensus sequences representing the A1 and A4 HPV16 sublineages from Ötzi and Ust’-Ishim, respectively, were generated from the mapping results using the Generate Consensus Sequence tool in Geneious Prime. Consensus sequences were built with a 60% highest-quality threshold, total quality assignment, and trimming to reference sequence, while positions with no coverage were designated as “?”. For each sublineage, the set of reads aligned to the corresponding reference genome was used to construct a consensus, reflecting the majority base at each position, weighted by read quality and coverage. This procedure synthesized the genomic information obtained from each mapping into a single sequence, reducing the impact of point variations and alignment artifacts [31]. The consensus sequences generated from Ötzi (A1) and Ust’-Ishim (A4) HPV16 sublineages are provided as **Supplementary Data 1 and 2**, respectively, in FASTA format.

Subsequently, two independent multiple sequence alignments were performed using the default parameters MAFFT v.7 [32] in Geneious Prime (www.geneious.com): one containing the five HPV16 sublineages reference sequences and the A1 consensus sequence derived from Ötzi, and another containing the five HPV16 sublineages reference sequences and the A4 consensus sequence derived from Ust’-Ishim. Generating separate alignments for each individual sample allowed a direct and controlled comparison of sublineage differences within each genetic context, avoiding cross-individual interference. MAFFT was chosen for its accuracy and performance in viral genome alignments, preserving positional homology necessary for downstream comparative analyses. From each multiple sequence alignment, pairwise distance matrices were calculated between the six sequences, quantifying the genetic divergence among the HPV16 sublineages for each individual [33]. The resulting distance tables allow quantitative assessment of similarity among the observed viral variants.

### Identification of Single-Nucleotide Variants in E6 and E7

To investigate potential nucleotide and amino acid substitutions in the E6 and E7 oncogenes, we analyzed the aligned BAM files generated from each ancient genome. For Ötzi, we used the BAM file corresponding to the mapping against the HPV16 A1 reference genome, whereas for Ust’-Ishim we used the BAM file obtained from the mapping against the HPV16 A4 reference, as these were the references with the highest mapping performance for each individual.

Single-nucleotide variants were detected using the Find Variations/SNPs tool (Geneious Prime (www.geneious.com). Given the limited and heterogeneous coverage typical of ancient DNA, all filters related to minimum coverage, minimum variant frequency, maximum variant *p*-value, and minimum strand-bias *p*-value were disabled. The options Only find SNVs and Merge adjacent variations were enabled to restrict the analysis to single-nucleotide substitutions and to avoid redundant calls. Variant calling was performed independently for each BAM file, restricted to the E6 and E7 coding regions. For every single nucleotide variant (SNV) identified by the software, the corresponding position in the reference sequences (A1 for Ötzi and A4 for Ust’) was manually annotated. We recorded (i) the observed nucleotides, and (ii) the resulting amino-acid substitution in the encoded proteins. These annotations were incorporated directly into the respective reference sequences to document all differences between the ancient reads and the HPV16 reference genomes used for mapping.

