## Supplementary material for "Oncogenic HPV types identified in Paleolithic and Chalcolithic human genome sequencing data from Ust’-Ishim and Ötzi": 03 Supplementary Table 1.pdf

**Supplementary Table 1.** Detailed metadata for sequencing runs. Data was retrieved from the NCBI SRA [23] and ENA [19].

| Sample / Individual | Run accession | Sequencing platform | Library layout | # Reads | Total data size (GB) | BioProject |
|---|---|---|---|---|---|---|
| Ötzi | ERR11837083 | Illumina | Paired-end | 132,746,967 | 2.1 | PRJEB56570 |
| Ötzi | ERR11837081 | Illumina | Paired-end | 678,475,568 | 10.3 | PRJEB56570 |
| Ötzi | ERR11837082 | Illumina | Paired-end | 148,108,402 | 2.2 | PRJEB56570 |
| Ötzi | ERR11837084 | Illumina | Paired-end | 202,111,622 | 3.5 | PRJEB56570 |
| Ötzi | ERR11837085 | Illumina | Paired-end | 224,865,862 | 3.8 | PRJEB56570 |
| Ötzi | ERR107307 | SOLiD | Paired-end | 475,180,823 | 25.6 | PRJEB2830 |
| Ötzi | ERR107308 | SOLiD | Paired-end | 517,633,616 | 29.4 | PRJEB2830 |
| Ötzi | ERR107309 | SOLiD | Paired-end | 526,112,461 | 28.8 | PRJEB2830 |
| Ötzi | ERR069107 | SOLiD | Paired-end | 251,115,149 | 9.0 | PRJEB2830 |
| Ötzi | ERR069108 | SOLiD | Paired-end | 296,268,307 | 10.1 | PRJEB2830 |
| Ötzi | ERR069109 | SOLiD | Paired-end | 259,438,904 | 9.1 | PRJEB2830 |
| Ust'-Ishim | ERR566093 | Illumina | Paired-end | 2,039,299,094 | 35.0 | PRJEB6622 |
| Total | — | — | — | 5,751,356,775 | 168.9 | — |
