## Supplementary material for "Oncogenic HPV types identified in Paleolithic and Chalcolithic human genome sequencing data from Ust’-Ishim and Ötzi": 32 Supplementary Data 31 FastQC ERR069107_reverse.html

ERR069107\_reverse.gz FastQC Report 

FastQC Report

Mon 24 Mar 2025  
ERR069107\_reverse.gz

### Summary

- Basic Statistics
- Per base sequence quality
- Per sequence quality scores
- Per base sequence content
- Per sequence GC content
- Per base N content
- Sequence Length Distribution
- Sequence Duplication Levels
- Overrepresented sequences
- Adapter Content

### Basic Statistics

| Measure | Value |
| --- | --- |
| Filename | ERR069107\_reverse.gz |
| File type | Colorspace converted to bases |
| Encoding | Sanger / Illumina 1.9 |
| Total Sequences | 134071453 |
| Total Bases | 3.3 Gbp |
| Sequences flagged as poor quality | 0 |
| Sequence length | 25 |
| %GC | 43 |

### Per base sequence quality

### Per sequence quality scores

### Per base sequence content

### Per sequence GC content

### Per base N content

### Sequence Length Distribution

### Sequence Duplication Levels

### Overrepresented sequences

| Sequence | Count | Percentage | Possible Source |
| --- | --- | --- | --- |
| AGAGAGAGAGAGAGAGAGAGAGAGA | 986312 | 0.7356614535981795 | No Hit |
| CACACACACACACACACACACACAC | 755800 | 0.5637292526396354 | No Hit |
| CTCTCTCTCTCTCTCTCTCTCTCTC | 497038 | 0.37072619776858834 | No Hit |
| TGTGTGTGTGTGTGTGTGTGTGTGT | 355141 | 0.2648893497111574 | No Hit |
| TCTCTCTCTCTCTCTCTCTCTCTCT | 290012 | 0.2163115215884175 | No Hit |
| GAGAGAGAGAGAGAGAGAGAGAGAG | 282116 | 0.21042212468600605 | No Hit |
| TGGAATGGAATGGAATGGAATGGAA | 181013 | 0.13501233554916423 | No Hit |

### Adapter Content

Produced by FastQC (version 0.12.1)
