## Supplementary material for "Oncogenic HPV types identified in Paleolithic and Chalcolithic human genome sequencing data from Ust’-Ishim and Ötzi": 34 Supplementary Data 33 FastQC ERR069108_reverse.html

ERR069108\_reverse.gz FastQC Report 

FastQC Report

Mon 24 Mar 2025  
ERR069108\_reverse.gz

### Summary

- Basic Statistics
- Per base sequence quality
- Per sequence quality scores
- Per base sequence content
- Per sequence GC content
- Per base N content
- Sequence Length Distribution
- Sequence Duplication Levels
- Overrepresented sequences
- Adapter Content

### Basic Statistics

| Measure | Value |
| --- | --- |
| Filename | ERR069108\_reverse.gz |
| File type | Conventional base calls |
| Encoding | Sanger / Illumina 1.9 |
| Total Sequences | 154461015 |
| Total Bases | 3.8 Gbp |
| Sequences flagged as poor quality | 0 |
| Sequence length | 25 |
| %GC | 44 |

### Per base sequence quality

### Per sequence quality scores

### Per base sequence content

### Per sequence GC content

### Per base N content

### Sequence Length Distribution

### Sequence Duplication Levels

### Overrepresented sequences

| Sequence | Count | Percentage | Possible Source |
| --- | --- | --- | --- |
| AGAGAGAGAGAGAGAGAGAGAGAGA | 1201839 | 0.7780856548171718 | No Hit |
| CACACACACACACACACACACACAC | 895268 | 0.5796077411507363 | No Hit |
| CTCTCTCTCTCTCTCTCTCTCTCTC | 660602 | 0.4276820270797781 | No Hit |
| TGTGTGTGTGTGTGTGTGTGTGTGT | 476832 | 0.3087070222864973 | No Hit |
| TCTCTCTCTCTCTCTCTCTCTCTCT | 377483 | 0.24438723259717024 | No Hit |
| GAGAGAGAGAGAGAGAGAGAGAGAG | 346179 | 0.22412063005024274 | No Hit |
| TGGAATGGAATGGAATGGAATGGAA | 209260 | 0.13547755075932916 | No Hit |

### Adapter Content

Produced by FastQC (version 0.12.1)
