## Supplementary material for "Oncogenic HPV types identified in Paleolithic and Chalcolithic human genome sequencing data from Ust’-Ishim and Ötzi": 36 Supplementary Data 35 FastQC ERR069109_reverse.html

ERR069109\_reverse.gz FastQC Report 

FastQC Report

Mon 24 Mar 2025  
ERR069109\_reverse.gz

### Summary

- Basic Statistics
- Per base sequence quality
- Per sequence quality scores
- Per base sequence content
- Per sequence GC content
- Per base N content
- Sequence Length Distribution
- Sequence Duplication Levels
- Overrepresented sequences
- Adapter Content

### Basic Statistics

| Measure | Value |
| --- | --- |
| Filename | ERR069109\_reverse.gz |
| File type | Colorspace converted to bases |
| Encoding | Sanger / Illumina 1.9 |
| Total Sequences | 140436833 |
| Total Bases | 3.5 Gbp |
| Sequences flagged as poor quality | 0 |
| Sequence length | 25 |
| %GC | 44 |

### Per base sequence quality

### Per sequence quality scores

### Per base sequence content

### Per sequence GC content

### Per base N content

### Sequence Length Distribution

### Sequence Duplication Levels

### Overrepresented sequences

| Sequence | Count | Percentage | Possible Source |
| --- | --- | --- | --- |
| AGAGAGAGAGAGAGAGAGAGAGAGA | 1050727 | 0.7481847728651072 | No Hit |
| CTCTCTCTCTCTCTCTCTCTCTCTC | 693599 | 0.4938868138674133 | No Hit |
| CACACACACACACACACACACACAC | 683080 | 0.48639661362913245 | No Hit |
| TGTGTGTGTGTGTGTGTGTGTGTGT | 487334 | 0.3470129520793167 | No Hit |
| TCTCTCTCTCTCTCTCTCTCTCTCT | 451884 | 0.32177028657432055 | No Hit |
| GAGAGAGAGAGAGAGAGAGAGAGAG | 301618 | 0.21477129151723323 | No Hit |
| TGGAATGGAATGGAATGGAATGGAA | 172337 | 0.12271495754963373 | No Hit |

### Adapter Content

Produced by FastQC (version 0.12.1)
