## Supplementary material for "Oncogenic HPV types identified in Paleolithic and Chalcolithic human genome sequencing data from Ust’-Ishim and Ötzi": 37 Supplementary Data 36 FastQC ERR107307_forward.html

ERR107307\_forward.gz FastQC Report 

FastQC Report

Tue 25 Mar 2025  
ERR107307\_forward.gz

### Summary

- Basic Statistics
- Per base sequence quality
- Per sequence quality scores
- Per base sequence content
- Per sequence GC content
- Per base N content
- Sequence Length Distribution
- Sequence Duplication Levels
- Overrepresented sequences
- Adapter Content

### Basic Statistics

| Measure | Value |
| --- | --- |
| Filename | ERR107307\_forward.gz |
| File type | Colorspace converted to bases |
| Encoding | Sanger / Illumina 1.9 |
| Total Sequences | 433933475 |
| Total Bases | 21.6 Gbp |
| Sequences flagged as poor quality | 0 |
| Sequence length | 50 |
| %GC | 44 |

### Per base sequence quality

### Per sequence quality scores

### Per base sequence content

### Per sequence GC content

### Per base N content

### Sequence Length Distribution

### Sequence Duplication Levels

### Overrepresented sequences

| Sequence | Count | Percentage | Possible Source |
| --- | --- | --- | --- |
| TGTGTGTGTGTGTGTGTGTGTGTGTGTGTGTGTGTGTGTGTGTGTGTGTG | 557001 | 0.12836091983915277 | No Hit |
| TCTCTCTCTCTCTCTCTCTCTCTCTCTCTCTCTCTCTCTCTCTCTCTCTC | 456185 | 0.10512786550979963 | No Hit |

### Adapter Content

Produced by FastQC (version 0.12.1)
