## Supplementary material for "Oncogenic HPV types identified in Paleolithic and Chalcolithic human genome sequencing data from Ust’-Ishim and Ötzi": 40 Supplementary Data 39 FastQC ERR107308_reverse.html

ERR107308\_reverse.gz FastQC Report 

FastQC Report

Tue 25 Mar 2025  
ERR107308\_reverse.gz

### Summary

- Basic Statistics
- Per base sequence quality
- Per sequence quality scores
- Per base sequence content
- Per sequence GC content
- Per base N content
- Sequence Length Distribution
- Sequence Duplication Levels
- Overrepresented sequences
- Adapter Content

### Basic Statistics

| Measure | Value |
| --- | --- |
| Filename | ERR107308\_reverse.gz |
| File type | Colorspace converted to bases |
| Encoding | Sanger / Illumina 1.9 |
| Total Sequences | 517633616 |
| Total Bases | 12.9 Gbp |
| Sequences flagged as poor quality | 0 |
| Sequence length | 25 |
| %GC | 48 |

### Per base sequence quality

### Per sequence quality scores

### Per base sequence content

### Per sequence GC content

### Per base N content

### Sequence Length Distribution

### Sequence Duplication Levels

### Overrepresented sequences

| Sequence | Count | Percentage | Possible Source |
| --- | --- | --- | --- |
| AGAGAGAGAGAGAGAGAGAGAGAGA | 1214833 | 0.23468974240652868 | No Hit |
| CACACACACACACACACACACACAC | 885720 | 0.17110944355669513 | No Hit |
| ATCACCGACTGCCCATAGAGATTCT | 703166 | 0.13584241406763659 | ABI Solid3 Adapter B (95% over 21bp) |
| CTCTCTCTCTCTCTCTCTCTCTCTC | 638224 | 0.12329647462463102 | No Hit |

### Adapter Content

Produced by FastQC (version 0.12.1)
