## Supplementary material for "Oncogenic HPV types identified in Paleolithic and Chalcolithic human genome sequencing data from Ust’-Ishim and Ötzi": 49 Supplementary Data 46 Duplicates.docx

After the initial simultaneous mapping with the five Human papillomavirus reference genomes, reads aligning to HPV16 (A1) were identified and extracted to enable an additional exploratory analysis. This subset was not used in the main analytical workflow but was examined separately to further investigate alignment behavior and potential lineage-specific affinities. From this HPV16-specific subset, duplicate removal was performed using the default settings of the Dedupe Duplicate Read Remover 38.84 tool in Geneious Prime [33] separately for the Ötzi (SOLiD) and Ust’-Ishim datasets. This step aimed to eliminate potential redundancies introduced during amplification or sequencing, enabling a more accurate assessment of genome coverage and read distribution along the viral genome.

To further assess the potential effect of data preprocessing on alignment outcomes, a complementary exploratory analysis was conducted in which the deduplicated read sets were remapped against the five reference genomes representing the HPV16 sublineages A1, A4, B1, C1, and D3. This comparison allowed evaluation of whether alignment affinities to specific sublineages persisted after removing potentially redundant reads.

Duplicate removal is generally avoided in ancient DNA research, particularly in viral paleogenomics, because identical reads may originate from genuinely independent molecules rather than PCR artifacts, and removing them can artificially reduce coverage estimates [37]. For this reason, the main analytical workflow retained duplicates, following standard practice for ancient viral datasets. However, to ensure that the HPV16 signal detected in the primary analysis was not driven by read redundancy, this supplementary deduplicated analysis was performed to test the robustness of the results.

To determine whether the mapping signal persisted after duplicate removal, real deduplicated mappings were compared to randomized mappings using Welch’s t-test, a statistical test appropriate for datasets with unequal variances and differing sample sizes. All HPV16 sublineages in the Ötzi dataset (A1, A4, B1, C1, and D3) showed real coverage values that were substantially higher than those obtained from randomized mappings, yielding extremely high *t* statistics and *p*-values approaching zero. These results demonstrate that the HPV16 signal in Ötzi remains significantly non-random even after removing duplicates, confirming that the positive alignment results observed in the main workflow are not artifacts caused by read redundancy.

For the Ust’-Ishim dataset, measurable coverage after deduplication was obtained only for the A1 and A4 references. The B1, C1, and D3 references did not produce contigs (“nd”), consistent with the extremely low viral read count in this individual. Even so, for A1 and A4, real coverage values still exceeded random expectations, indicating that a detectable, although weak, HPV16-related signal persists even under the more conservative deduplicated conditions.

Supplementary Table S1 summarizes the results of these comparisons. The table reports the mean coverage, standard deviation, and number of aligned reads for the real and randomized mappings, along with the corresponding Welch’s *t* statistics, critical values, and p-values. Cases marked “nd” indicate that no contigs were generated, preventing calculation of coverage metrics and statistical testing.

Taken together, these supplementary analyses confirm that the HPV16 signal detected in both individuals is not dependent on the retention of duplicate reads. The positive alignment pattern remains detectable and statistically robust even after duplicate removal, demonstrating that the findings of the main workflow are not driven by PCR artifacts or sequencing redundancies. Retaining duplicates in the primary analytical framework is therefore methodologically appropriate for ancient viral DNA and is further validated by the consistent results obtained under deduplicated conditions.

**Supplementary Table S1.** Results of the comparative analysis between real and randomized mappings after duplicate removal for the Ötzi (SOLiD) and Ust’-Ishim datasets. For each HPV16 reference genome (A1, A4, B1, C1, and D3), the table reports the mean genome coverage, standard deviation, and number of aligned reads obtained from the real deduplicated mappings, alongside the corresponding values from 100 randomized mapping iterations. Welch’s t-test statistics, critical values, and p-values are provided to assess whether real mappings differ significantly from random expectations. “nd” indicates that no contigs were generated, preventing calculation of coverage metrics and statistical testing.

|  | **Real** | | | **Random** | | | **Results** | | |
| --- | --- | --- | --- | --- | --- | --- | --- | --- | --- |
|  | Coverage Mean | Coverage SD | # Reads | Coverage Mean | Coverage SD | # Reads | *t* | Critical Value | *p*-value |
| Ötzi |  |  |  |  |  |  |  |  |  |
| A1 | 1587.6 | 8813.3 | 71,038 | 0.1 | 0.8 | 18 | 48.008 | 1.6449 | 0 |
| A4 | 677.8 | 4099.5 | 36,353 | 0.1 | 0.8 | 18 | 31.5181 | 1.6449 | 0 |
| B1 | 17.2 | 99.4 | 2,821 | 0.1 | 0.8 | 18 | 9.0911 | 1.6454 | 0 |
| C1 | 7.4 | 24.0 | 1,751 | 0.1 | 0.8 | 18 | 12.0912 | 1.6464 | 0 |
| D3 | 6.6 | 19.3 | 1,495 | 0.1 | 0.8 | 18 | 12.1818 | 1.647 | 0 |
| Ust-Ishim |  |  |  |  |  |  |  |  |  |
| A1 | 474.6 | 2223.9 | 12,862 | nd | nd | nd | nd | nd | nd |
| A4 | 279.1 | 1363.3 | 8,139 | nd | nd | nd | nd | nd | nd |
| B1 | 0.0 | 0.3 | 8 | nd | nd | nd | nd | nd | nd |
| C1 | 0.0 | 0.3 | 9 | nd | nd | nd | nd | nd | nd |
| D3 | 0.0 | 0.3 | 8 | nd | nd | nd | nd | nd | nd |
