## Supplementary figures and images for "Oncogenic HPV types identified in Paleolithic and Chalcolithic human genome sequencing data from Ust’-Ishim and Ötzi"

### 04 Supplementary Data 3 ERR107307 Run Browser SRA Archive NCBI.png

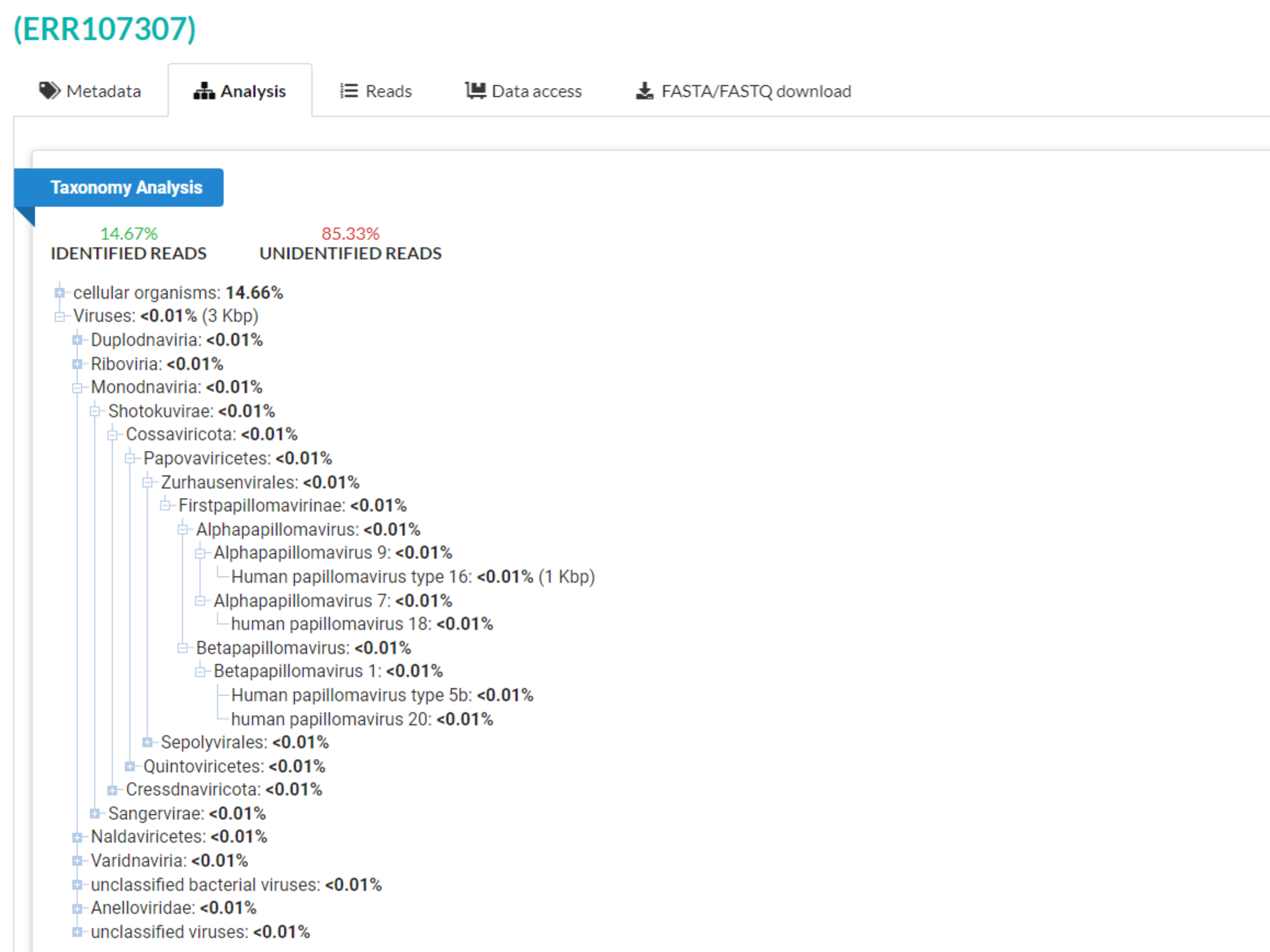

### 05 Supplementary Data 4 ERR107308 Run Browser SRA Archive NCBI.png

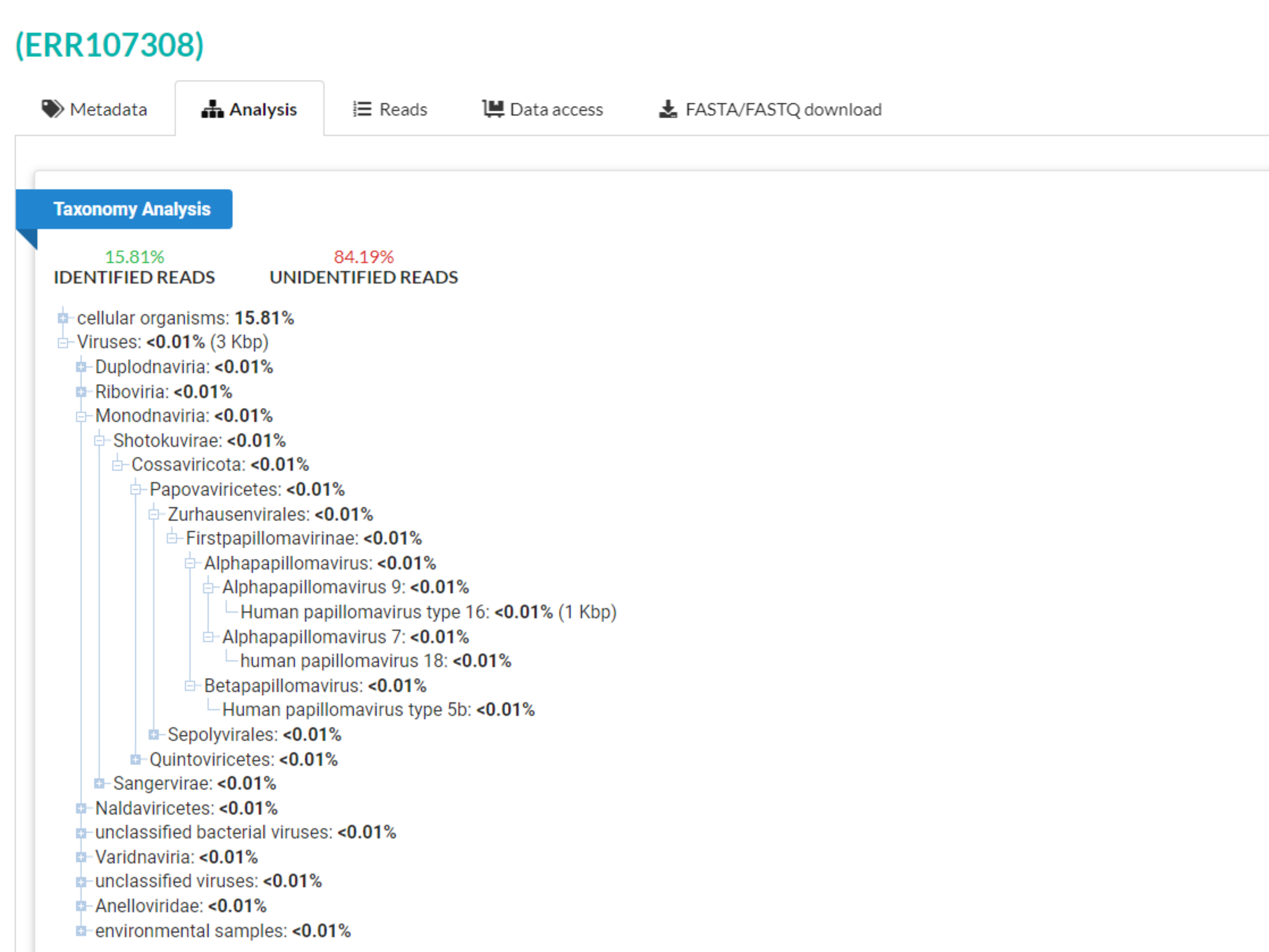

### 06 Supplementary Data 5 ERR107309 Run Browser SRA Archive NCBI.png

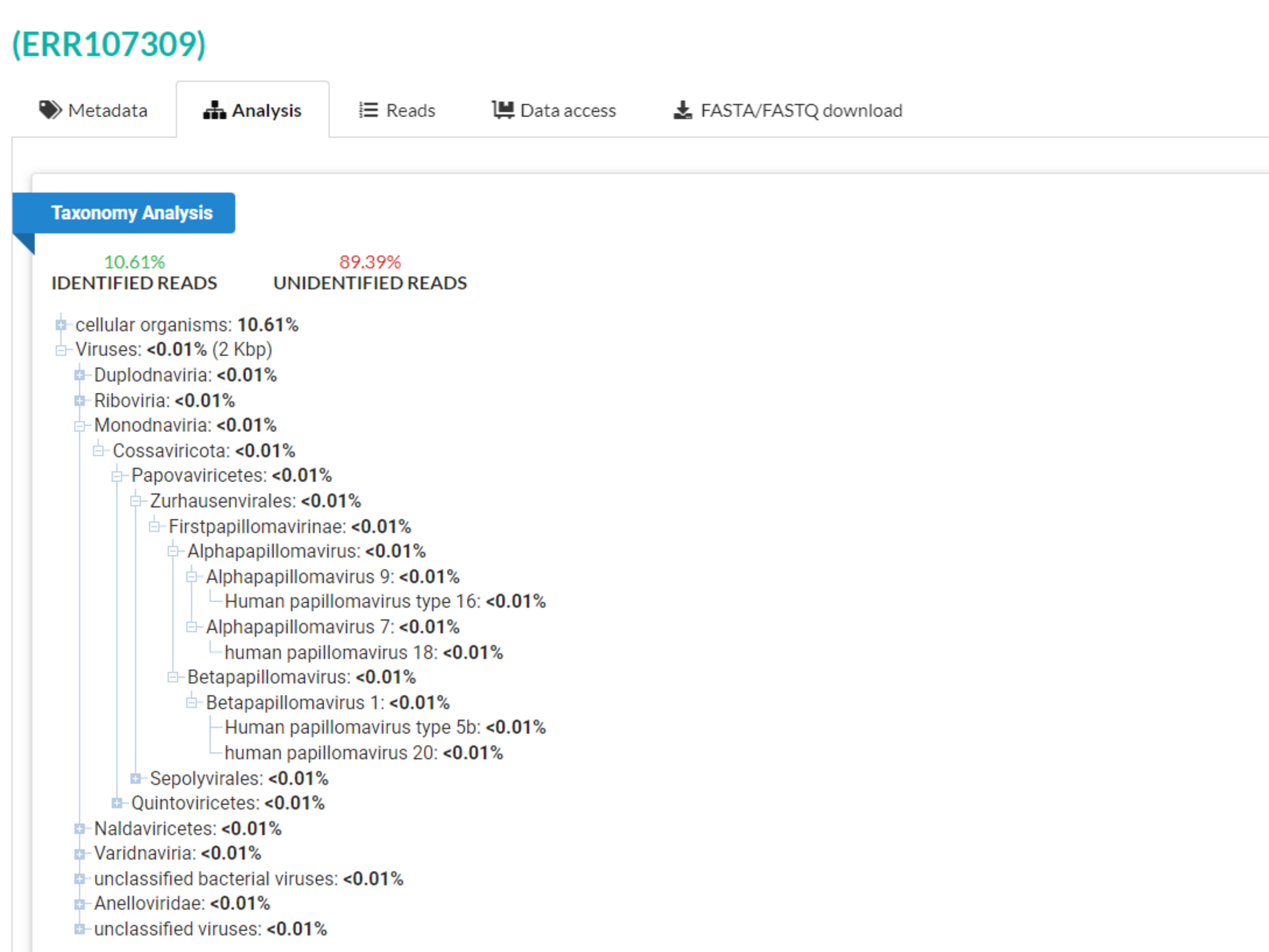

### 07 Supplementary Data 6 ERR11837082 Run Browser SRA Archive NCBI.png

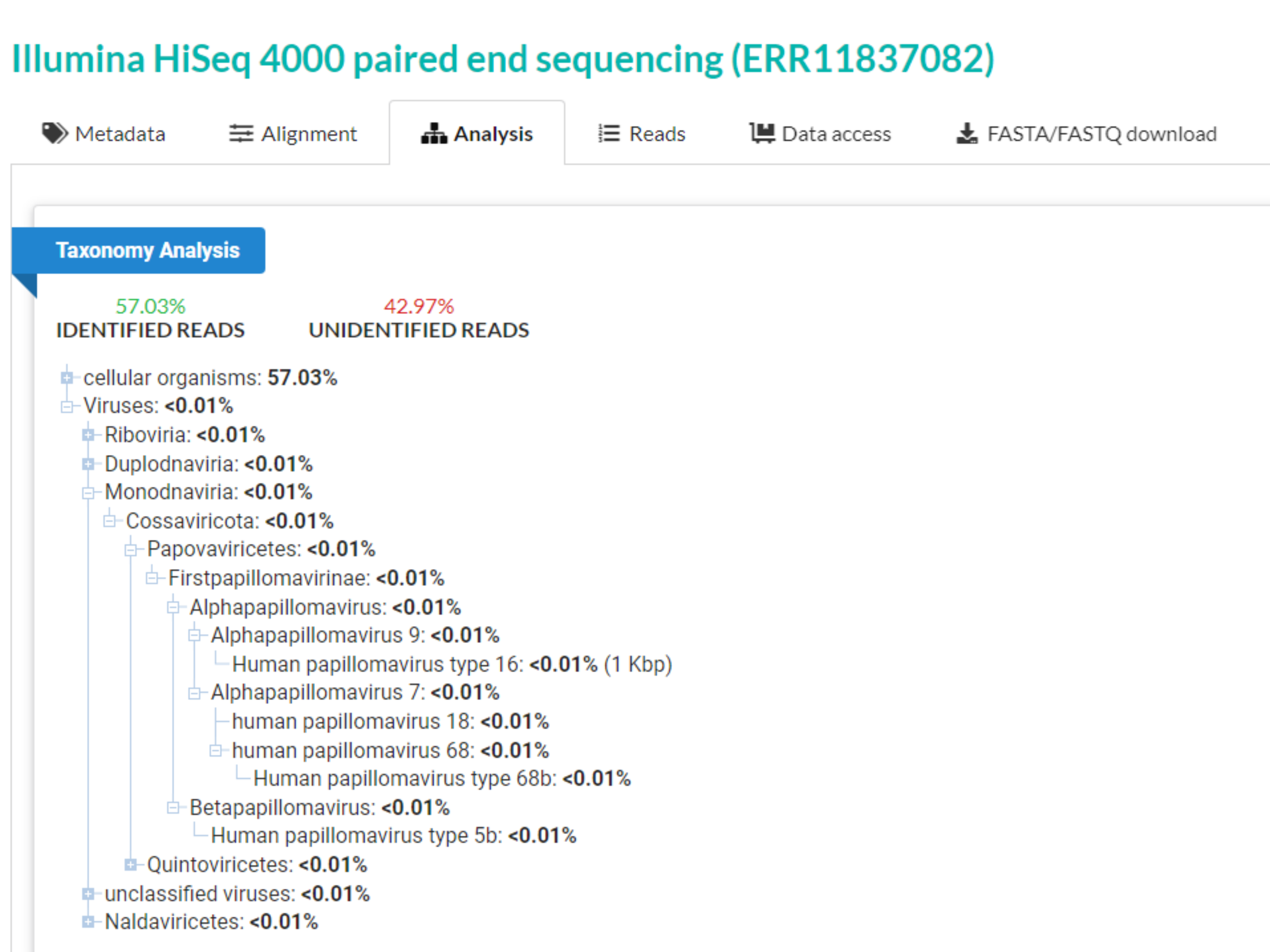

### 08 Supplementary Data 7 ERR11837083 Run Browser SRA Archive NCBI.png

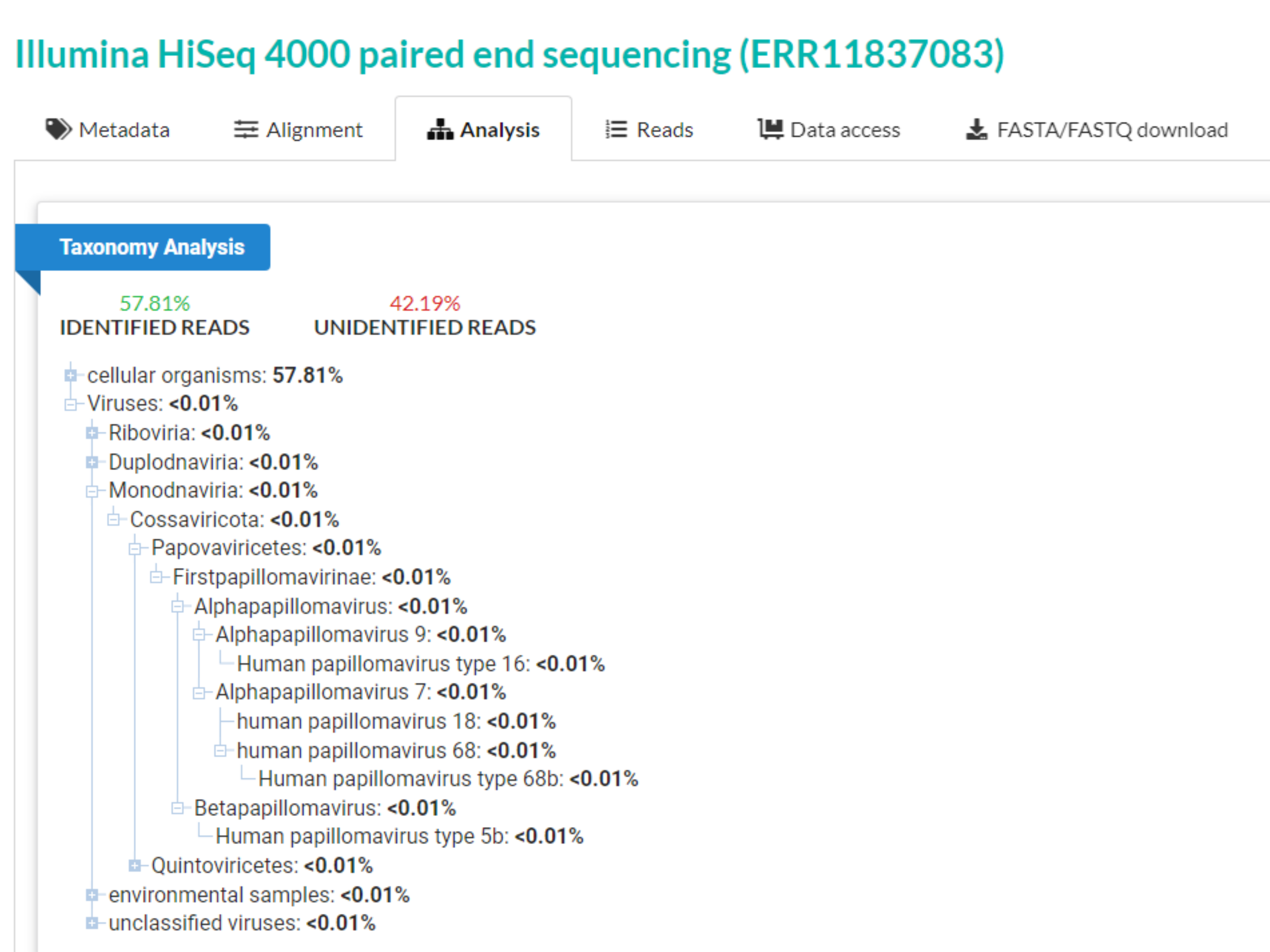

### 09 Supplementary Data 8 ERR11837084 Run Browser SRA Archive NCBI.png

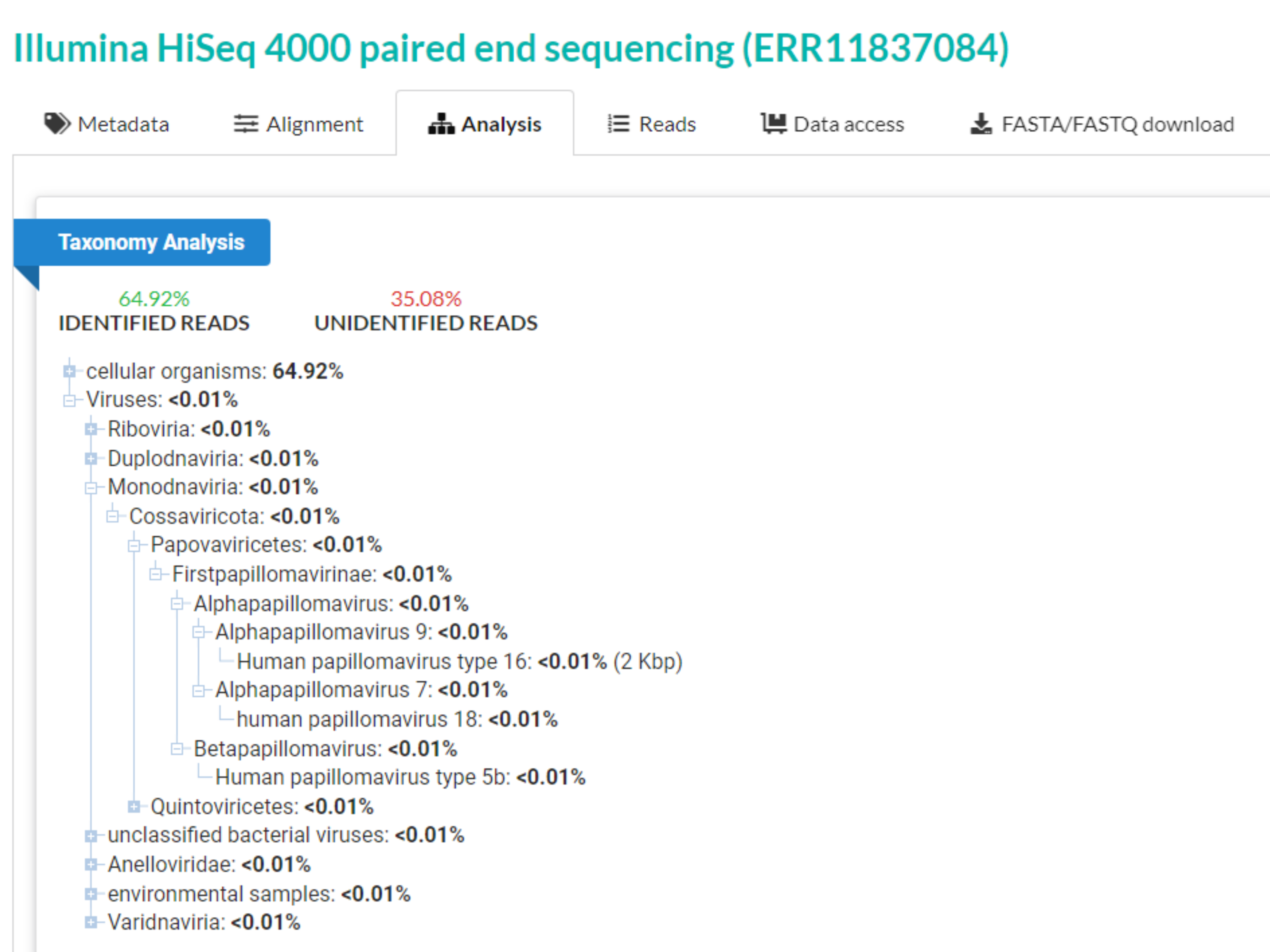

### 10 Supplementary Data 9 ERR11837085 Run Browser SRA Archive NCBI.png

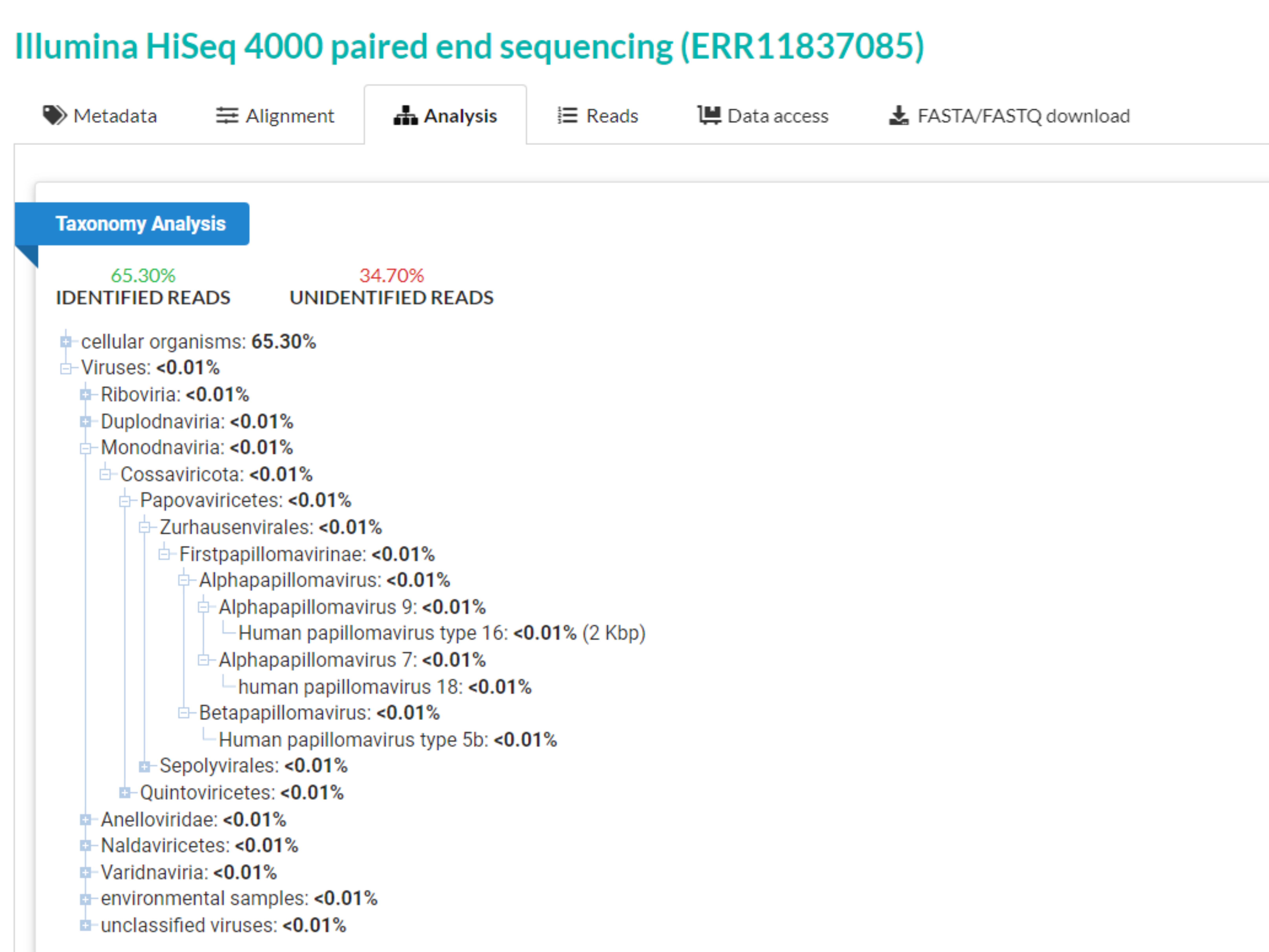
